# Characterisation of ovine bone marrow-derived stromal cells (oBMSC) and evaluation of chondrogenically induced *micro*-pellets for cartilage tissue repair *in vivo*

**DOI:** 10.1101/2020.05.15.094847

**Authors:** K. Futrega, E. Music, P.G. Robey, S. Gronthos, R.W. Crawford, S. Saifzadeh, T.J. Klein, M.R. Doran

**Author notes:** co-first author (K. Futrega, E. Music). **Corresponding Author Dr. Michael R. Doran**, School of Biomedical Sciences, Institute of Health and Biomedical Innovation, Queensland University of Technology, Translational Research Institute, 37 Kent Street, Brisbane, Queensland, Australia 4102, or **Dr. Michael R. Doran**, Skeletal Biology Section, National Institute of Dental and Craniofacial, Research, National Institutes of Health, Building 30, 30 Convent Dr MSC 4320, Bethesda, MD 20892-4320, United States.

## Abstract

**Background:** Bone marrow stromal cells (BMSC) show promise in cartilage repair, and sheep are the most common large animal pre-clinical model. The objective of this study was to characterize ovine BMSC (oBMSC) *in vitro*, and to evaluate the capacity of chondrogenic *micro*-pellets manufactured from oBMSC or ovine articular chondrocytes (oACh) to repair osteochondral defects in sheep.

**Methods:** oBMSC were characterised for surface marker expression using flow cytometry and evaluated for tri-lineage differentiation. oBMSC *micro*-pellets were manufactured in a microwell platform, and chondrogenesis was compared at 2%, 5%, and 20% O_2_. The capacity of cartilage *micro*-pellets manufactured from oBMSC or oACh to repair osteochondral defects in adult sheep was evaluated in an 8-week pilot study. Expanded oBMSC were positive for CD44 and CD146 and negative for CD45.

**Results:** The common adipogenic induction medium ingredient, 3-Isobutyl-1-methylxanthine (IBMX) was toxic to oBMSC, but adipogenesis could be restored by excluding IBMX from the medium. BMSC chondrogenesis was optimal in a 2% O_2_ atmosphere. *Micro*-pellets formed from oBMSC or oACh appeared morphologically similar, but hypertrophic genes were elevated in oBMSC *micro*-pellets. While oACh *micro*-pellets formed cartilage-like repair tissue in sheep, oBMSC *micro*-pellets did not.

**Conclusion:** The sensitivity of oBMSC to IBMX highlights species-species differences between oBMSC and hBMSC. *Micro*-pellets manufactured from oBMSC were not effective in repairing osteochondral defects, while oACh *micro*-pellets enabled modest repair. While oBMSC can be driven to form cartilage-like tissue in vitro, their effective use in cartilage repair will require mitigation of hypertrophy.

## Introduction

Cartilage has a limited capacity for regeneration. A number of therapies that use expanded autologous chondrocytes have been approved for the repair of focal cartilage defects, demonstrating the potential and promise of cell-based repair strategies [1]. While efficacious, these therapies require the harvest of autologous cartilage to source chondrocytes, and this limits the cell number available for input into these repair processes, and also causes donor site morbidity. This limitation has motivated investment into strategies that utilise bone marrow-derived stromal cells (BMSC, sometimes referred to as “*mesenchymal stem cells*”) as an alternative input cell source based on their chondrogenic potential. Thus far, no BMSC-based cartilage repair therapies have successfully passed the regulatory and efficacy hurdles required for approval [1]. A major efficacy limitation for BMSC-derived engineered cartilage is the observed hypertrophic conversion of these cells when transplanted *in vivo* in mice, resulting in remodelled, mineralized bone-like tissue [2, 3].

The majority of pre-clinical experimentation is performed in small animal models such as mice, rats, or rabbits[4, 5]. While some orthopaedic studies are performed using the joints of these small animals as models, these joints are anatomically different and experience reduced mechanical forces relative to human joints, and their cartilage can often heal spontaneously [4], unlike that observed in humans. In some studies using immunocompromised small animals, human tissue is implanted at ectopic sites, typically in subcutaneous pouches on the backs of mice [2, 3]. While this allows for the transplantation of human cells, the implant sites are dissimilar to human joints, as they do not provide mechanical load, and they are more vascular than joint capsules. Large animal models such as sheep, pigs, goats, or horses are more appropriate for orthopaedic studies due to their joint anatomy, and healing properties being more similar to that of humans [6]. However, unlike immunocompromised small animal models, these immune competent large animal models require that species-matched cells be used. Therefore, large animal models need to be characterised independently to determine their suitability as models for cellular therapies. While sheep are the most commonly used large animal models for the study of cartilage repair [6], the biology of sheep or ovine BMSC (oBMSC) remains poorly characterised relative to human BMSC (hBMSC).

Our team has developed *micro*-pellet models to characterise and optimise *in vitro* hBMSC chondrogenesis [3, 7, 8]. Pellet models for studying BMSC chondrogenesis were first developed by Johnstone *et al*. in 1998 [9]. While the traditional pellet model mimics aspects of developmental mesenchymal condensation, these pellets, formed from ~2×10^5^ cells each, have a large diameter, and suffer steep diffusion gradients and radial tissue heterogeneity [8]. By contrast, *micro*-pellets can be formed from fewer cells (5×10^3^ each), resulting in reduced diffusion gradients and more homogeneous tissue. Hereafter, traditional pellet cultures assembled from several hundred thousand cells will be referred to as *macro*-pellets, while smaller pellets will be referred to as *micro*-pellets. Previous data demonstrate that *micro*-pellets enable characterisation and optimisation of cell culture variables that are obfuscated by the radial heterogeneity suffered by traditional *macro*-pellet models [7]. In addition to being an excellent tissue culture model, *micro*-pellets could theoretically be used as building blocks to fill cartilage defects and facilitate repair [3, 10]. To expedite high-throughput *micro*-pellet manufacture, we developed a microwell culture platform called the *Microwell-mesh [3]*. The *Microwell-mesh* has a nylon mesh bonded over an array of microwells. The mesh openings are large enough to allow a single cell suspension to be centrifuged through the mesh, and into the microwells. Within the first three hours of culture, cells self-assemble into *micro*-pellets, becoming too large to escape back through the mesh; thus, they are retained in discrete microwells during weeks of culture manipulation. This feature of the *Microwell-mesh* allows hundreds to thousands of *micro*-pellets to be efficiently cultured simultaneously, facilitating large scale experimentation with *micro*-pellets that would otherwise be tedious or costly.

A low-oxygen atmosphere has been shown to enhance hBMSC chondrogenesis in cell culture [8], but there are few corresponding studies that optimise this crucial parameter for oBMSC. Oxygen concentration directly influences BMSC differentiation through hypoxia-inducible factors (HIFs) [8, 11–13]. HIF-1α is a key mediator of enhanced chondrogenesis under low oxygen conditions [14]. It upregulates the expression of chondrogenic genes, including *SOX9*, and downregulates genes associated with osteogenesis and hypertrophy, such as *COL1A1* and *RUNX2* [13, 15, 16]. Accurate evaluation of oxygen culture conditions is dependent on efficient and reproducible oxygen transport across the engineered oBMSC cartilage-like tissue. Compared with traditional *macro*-pellets, the smaller diameter *micro*-pellets exhibited a reduced diffusion gradient, yielding more homogeneous tissue, which allowed for the reliable optimisation of oxygen culture conditions during hBMSC chondrogenesis [3].

In this study, we aimed to optimise the chondrogenic culture conditions for oBMSC using the *micro*-pellet model, and to evaluate the use of *micro*-pellets as building blocks to repair osteochondral defects in sheep. We used the *Microwell-mesh* platform to manufacture *micro*-pellets formed from 5,000 oBMSC each and compared them to traditional *macro*-pellets manufactured from 200,000 oBMSC each. Chondrogenic induction was performed in atmospheres containing 2%, 5%, or 20% O_2_. The resulting cartilage-like tissue was evaluated based on glycosaminoglycan (GAG) accumulation, histological analysis, and gene expression. A pilot study was then performed to evaluate the capacity of autologous *micro*-pellets, derived from either oBMSC or expanded ovine articular chondrocytes (oACh), to repair osteochondral defects formed in the stifle joint of adult Merino sheep. Two sheep received oBMSC *micro*-pellets, while another two sheep received oACh *micro*-pellets. Each sheep had three replicate defects, 6 mm in diameter and 1.5 mm deep. Engineered constructs were assembled during the surgery using a mold to cast a 6 mm diameter by 1.5 mm deep layer of *micro*-pellets, set with Baxter Tisseel fibrin glue, on a CelGro™ collagen scaffold, and fixed in place with sutures. The pilot study was terminated 8 weeks later, and defect infill was characterised via histology.

## Materials and Methods

### Animal studies

All procedures were approved by the Queensland University of Technology (QUT) Animal Ethics Committee (approval number 1500001091). For surgical knee procedures, analgesia was provided just prior to surgery with slow release transdermal fentanyl patches (2-3 mg/kg/h). A first patch was applied just prior to surgery and two more patches were applied in 3-day intervals post-surgery. Trisoprim (1 mL/30 kg/day) was administered postoperatively for two days to prevent bacterial infection. For euthanasia, sheep were injected in the external jugular with 100 mg/kg Pentobarbital sodium.

### Bone marrow aspiration and BMSC culture

Bone marrow aspiration from the iliac crest of sheep was performed as described previously [17]. Briefly, an 11-guage Jamshidi needle was used to aspirate ~30 mL of bone marrow from the iliac crest into a 35 mL syringe containing 5 mL of heparin (1000 IU/mL). Aspirates were diluted in cell culture medium at a ratio of 5 mL aspirate to 25 mL medium, consisting of low glucose Dulbecco’s Modified Eagle’s Medium (LG-DMEM; Thermo Fisher Scientific) supplemented with 10% fetal bovine serum (FBS; Thermo Fisher Scientific), 10 ng/mL fibroblast growth factor-1 (FGF-1; Peprotech), 5 <g/mL porcine heparin sodium salt (Sigma-Aldrich), and 100 U/mL penicillin/streptomycin (PenStrep; Thermo Fisher Scientific). Cells were seeded into Nunc™ T175 cm^2^ tissue culture flasks (Thermo Fisher Scientific) and incubated overnight in a normoxic incubator (20% O_2_) with 5% CO_2_ at 37**°**C. Following overnight incubation, media was exchanged to remove non-adherent cells, and cultures were moved to a hypoxic incubator (2% O_2_, 5% CO_2_). Cells were cultured until 80% confluency, enzymatically harvested (0.25% Trypsin/EDTA; Thermo Fisher Scientific), and reseeded at 1,500 cells/cm^2^ in new T175 flasks. Culture medium was exchanged twice weekly. Cells were cryopreserved at low passage (P0-P2) in 90% FBS and 10% DMSO (Sigma-Aldrich).

hBMSC isolation and expansion was performed as previously described [5], and cultured as described for oBMSC. Ethics approval was granted by the Mater Health Services Human Research Ethics Committee (Ethics number: 1541A).

### Flow cytometry characterization

oBMSC were stained with the following anti-human antibodies: CD34-APC, CD90-FITC, CD73-APC, CD105-PE, CD146-APC, CD271-PE, HLA-DR-PE, and isotype controls (Miltenyi Biotec) and anti-sheep antibodies: CD45-FITC and CD44-FITC (Bio-Rad). Cells were stained as per the manufacturers’ instructions and analysis was completed using an LSR II flow cytometer (BD Biosciences) and FlowJo software (BD). Analysis included hBMSC, and sheep bone marrow mononuclear cells (MNC) where haematopoietic cells (CD45^+^) would be expected.

### Articular cartilage biopsy and oACh culture

Two oACh donors (oACh 1 and oACh 2) were used in cartilage defect repair component of this study. Articular cartilage biopsies were harvested from the femoral trochlear grooves. The knee joint capsule was accessed via a medial parapatellar incision (~5 cm). The patella was displaced laterally and retained in a proximal position with small wound retractors. The joint capsule was opened to expose the femoral trochlear groove. Five thin (~1 mm thick and 4-6 mm wide) shavings of articular cartilage were aseptically removed with a scalpel and transferred to a sterile specimen jar containing LG-DMEM with PenStrep and 1X Antibiotic-Antimycotic (Anti-Anti; Thermo Fisher Scientific). The biopsied tissues were chopped up coarsely, and resuspended in 10 mL of digestion solution containing type II clostridial collagenase (~430 U/mL; Sigma-Aldrich), PenStrep, and Antibiotic-Antimicotic (Gibco), and incubated overnight in a normoxic incubator (20% O_2_) with 5% CO_2_ at 37**°**C. Following digestion, the cells were washed twice by centrifugation (500 x g) with LG-DMEM containing 10% FBS. oACh were grown in a hypoxic (2% O_2_) incubator in growth medium as described above for oBMSC.

### *Fabrication of the* Microwell-mesh *culture device*

The *Microwell-mesh* was prepared as detailed in a previous paper published by our laboratory [3]. Briefly, a ~4 mm layer of polydimethylsiloxane (PDMS; Dow Corning) was poured into a polystyrene negative template that had an inverted microwell pattern. PDMS was cured at 80°C for 60 minutes. A wad punch was used to create discs that fit snuggly into Nunc™ 6-well plates (Thermo Fisher Scientific). Individual microwells measured 2 mm x 2 mm with a depth of 0.8 mm [3]. A nylon mesh (36 μm square pore openings; part number CMN-0035, Amazon.com) was bonded to the open face of the PDMS discs with silicone glue (Selleys, Aquarium Safe). Plates were sterilised by submerging in a 70% ethanol solution for a minimum of 30 minutes and rinsed 3 times with phosphate buffered saline (PBS; Thermo Fisher Scientific). Prior to cell seeding, microwells were rinsed with a sterile 5% Pluronic F-127 solution (Sigma-Aldrich) in PBS to render the PDMS surface non-adhesive and to promote cell aggregation [18, 19]. Wells were rinsed 3 times with PBS to remove excess Pluronic and then cells were seeded as described below.

### Osteogenic and adipogenic induction

To induce osteogenesis and adipogenesis, oBMSC were seeded at 30,000 cells/cm^2^ in standard tissue culture well plates and cultured for 21 days. Osteogenic induction medium was composed of 10 mM β-glycerol phosphate, 100 nM dexamethasone (Dex), 50 μM L-ascorbic acid 2-phosphate (all from Sigma-Aldrich), 10% FBS, and PenStrep in high glucose (HG)-DMEM (Thermo Fisher Scientific). Conventional adipogenic induction medium was composed of 10 <g/mL insulin, 100 nM Dex, 25 mM indomethacin, and 3-Isobutyl-1-methylxanthine (IBMX) (all from Sigma-Aldrich), 10% FBS, and PenStrep in HG-DMEM. We noticed that adipogenic induction was not successful using conventional adipogenic induction medium, which appeared to be toxic. We then tested whether a single ingredient could be removed from the adipogenic induction medium to eliminate the toxicity and demonstrate adipogenic induction. We included a hBMSC population in this adipogenic assay for comparison.

To assess osteogenic and adipogenic induction, monolayers were fixed for 20 minutes in 4% paraformaldehyde (PFA), then washed and stained. Mineralised matrix deposition was assessed in osteogenic cultures using Alizarin Red S staining (Sigma-Aldrich). Lipid vacuoles in adipogenic cultures were stained using Oil Red O (Sigma-Aldrich). Wells were rinsed with distilled water, incubated with Alizarin Red S or Oil Red O stain for 10 minutes, then washed with water and visualised.

### Chondrogenic induction cultures

To induce chondrogenesis, cells were resuspended in chondrogenic medium composed of HG-DMEM, PenStrep, GlutaMax, 1X ITS-X, 100 μM sodium pyruvate (all from Thermo Fisher Scientific), 10 ng/mL TGF-β1 (PeproTech), 100 nM Dex, 200 μM ascorbic acid 2-phosphate, and 40 μg/mL L-proline (Sigma-Aldrich). Prior to cell seeding, 3 mL of cell-free chondrogenic induction medium was added to each well, and plates were centrifuged for 5 min at 2000 *xg* to ensure elimination of air bubbles from microwells. To generate *micro*-pellets, each well was seeded with 1.2×10^6^ oBMSC in 1 mL of chondrogenic medium and the plate was centrifuged at 500 *xg* for 3 minutes to force the cells through the nylon mesh and into the microwells. The microwell array has ~250 microwells per well, and therefore, ~5,000 cells are seeded per *micro*-pellet. At culture harvest, the nylon mesh was peeled off of the PDMS microwell insert to liberate *micro*-pellets. Control *macro*-pellet cultures were established by seeding 2×10^5^ oBMSC in 1 mL of induction medium in 96-well, deep V-bottom plates (Corning). Cultures were maintained at 2%, 5%, or 20% O_2_, and 5% CO_2_ in a 37°C incubator for optimisation experiments, where indicated. Media was exchanged every second day.

### Quantification of glycosaminoglycans (GAG) and DNA

Tissues were incubated in an overnight papain digest (1.6 U/mL papain, 10 mM L-cysteine; both from Sigma-Aldrich) in a 60°C water bath. The 1,9-dimethylmethylene blue (DMMB, Sigma-Aldrich) assay was used to quantify GAG in the digested tissues [8]. Chondroitin sulfate from shark cartilage (Sigma-Aldrich) was used to generate a standard curve. DNA content in the tissues was quantified using the Quant-iT™ PicoGreen® dsDNA assay kit (Thermo Fisher Scientific) as per the manufacturer’s protocol.

### Quantitative PCR (qPCR)

At harvest, tissues were collected and stored in 350 μL RLT buffer (Qiagen) containing β-mercaptoethanol (Sigma-Aldrich) at −80°C. The samples were crushed in 1.5 mL microcentrifuge tubes using sterile micropestles, then RNA was isolated using the RNeasy Mini Kit (Qiagen) with on-column DNase I (Qiagen) digestion, as per manufacturer’s instructions. RNA was quantified with a NanoDrop Lite spectrophotometer (Thermo Fisher Scientific) and RNA was reverse-transcribed using the SuperScript III First-Strand Synthesis System for RT-PCR (Thermo Fisher Scientific). The master mix included 2X SYBR Green PCR Master Mix (Applied Biosystems), 200 nM of the forward and reverse primers, RNase-free water, and 1 μl of sample cDNA. The 5 μl reactions were run in triplicate in a 384-well plate inside a Viia7 Real Time PCR System (Applied Biosystems). The initial cycle was 50°C for 2 minutes and 95°C for 10 minutes, followed by 40 cycles of 95°C for 15 seconds and 60°C for 1 minute. The melt curve and electrophoretic gels were evaluated to confirm the specificity of products. Primer set information for genes of interest is given in Table 1. Gene expression values were normalised to GAPDH and calculated using the ΔCt method.

**Table 1.**
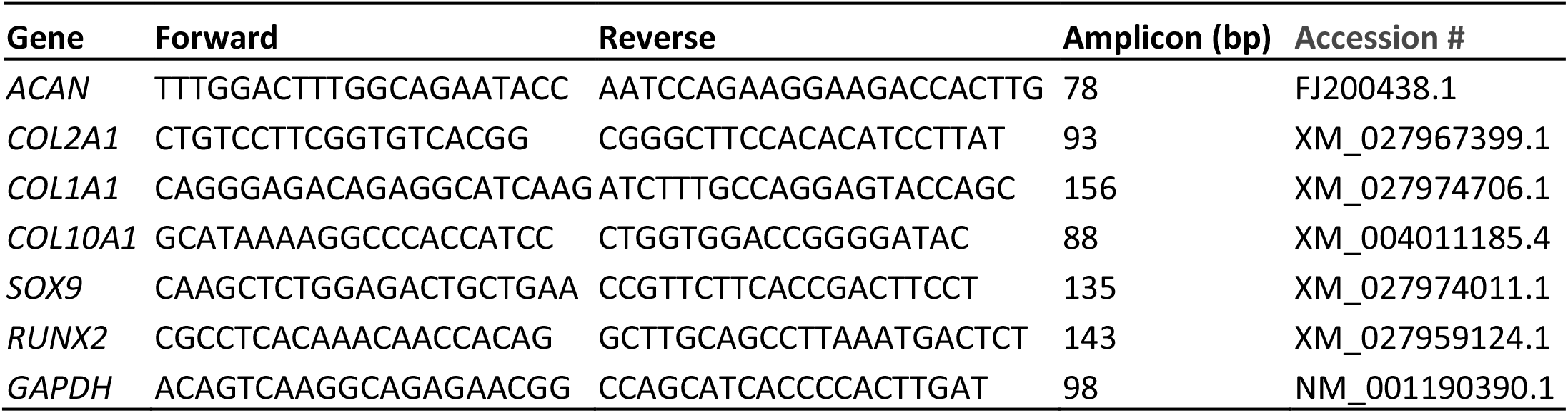
Primers used for qPCR for ovine genes.

### Histology and immunohistochemistry

Tissues were washed in PBS and fixed in 4% PFA for 20 min and frozen in Tissue-Tek OCT compound (Sakura Finetek). Samples were cryosectioned at 7 μm and collected onto poly-lysine coated slides, and then frozen until further processing. The sectioned tissues were fixed for 15 min with 4% PFA and washed with PBS. Sections were stained with Alcian blue (Sigma-Aldrich) for 30 min to visualise GAG distribution and counterstained with Nuclear Fast Red for 5 min. For immunohistochemical staining, sections were treated with hyaluronidase (2 U/mL, Sigma-Aldrich) for 30 min at 37°C. Slides were washed twice with 0.025% Triton X-100/PBS, then blocked for 60 min at room temperature with 10% normal goat serum (Thermo Fisher Scientific). Primary antibodies (all from Abcam) raised against collagen type I (1:800; ab6308), type II (1:100; ab34712), and type X (1:100; ab58632) were diluted in 1% BSA/PBS and incubated on sections at 4°C, overnight. Sections were rinsed twice for 5 min with 0.025% Triton X-100/PBS. Slides were incubated with 0.3% H_2_O_2_ for 15 min and rinsed twice with PBS. Sections were incubated for 60 min at room temperature with secondary antibodies, goat anti-rabbit (HRP; ab6721) or goat anti-mouse (HRP; ab97023), at 1:1000 (Abcam) in 1% BSA/PBS, then washed twice with PBS. A DAB chromogen kit (Abcam) was used to develop the signal for 8 minutes, then slides were rinsed in water, mounted, and coverslipped.

### *Autologous repair of* in vivo *cartilage defects in sheep*

We evaluated oBMSC *micro*-pellets and oACh *micro*-pellets *in vivo,* in sheep cartilage defects. Due to the high cost of sheep and the complexity of the procedure, we performed a pilot study with 2 sheep for oBMSC and 2 sheep for oACh *micro*-pellet evaluation. The procedure was carried out in two steps. In a first procedure, bone marrow aspirates or articular cartilage were collected from 2 sheep each (total of 4 sheep). The cells were isolated, expanded, and frozen at P0, as described above. In a second procedure, cells were thawed, expanded for one additional passage, induced for 10 days in *Microwell-mesh* cultures and implanted in cartilage defects created in the trochlear groove of knees of the same sheep from which the original cells were sourced (autologous implantation). All cell culture was carried out in an incubator set at 2% O_2_, 5% CO_2_, and 37°C. Sheep that received oACh *micro*-pellets received the implanted tissue in the knee opposite to the one where the original cartilage biopsy had been collected. Some *micro*-pellets were saved for GAG staining and qPCR analysis. Sheep were anaesthetised and prepared for surgery. Following exposure of the femoral trochlear groove, three 6 mm defects were made in the trochlear groove using a 6 mm biopsy punch, and the articular and calcified cartilage was removed using a curette until the subchondral bone was exposed (~1.5 mm deep). Approximately 5 mm of intact cartilage was left between defects. The defect sites and exposed cartilage were irrigated with saline regularly to clean and maintain tissue moisture.

To prepare the cartilage constructs, we used custom well chambers that consisted of a glass slide and a cylindric well made from a PDMS ring that had an internal diameter of 6 mm and 1.5 mm depth. The well chambers were sterilised by autoclaving and handled aseptically. A 6 mm wide disc was punched from a sheet of CelGro™ (Orthocell Ltd, Perth, Australia) collagen scaffold and placed in the bottom of the well. A dropper was used to drop *micro*-pellets into the well on top of the CelGro scaffold. Excess liquid was removed from the chamber with a pipette tip. Once the well was adequately filled with *micro*-pellets, two drops of fibrinogen were applied, followed by two drops of thrombin, both from a Baxter Tisseel fibrin glue kit. The gelled construct was removed from the chamber using forceps and inserted into the cartilage defect with the *micro-*pellets facing the subchondral bone and the CelGro scaffold sitting on top, adjacent to the superficial cartilage layer. To provide a scaffold-only control, one defect site in both the oBMSC and oACh sheep groups was filled with fibrin glue and the Celgro scaffold, but no cells. Four degradable sutures were used to fix the membrane to adjacent cartilage tissue. The patella was returned to its natural position and the incision was sutured to close the knee joint. Knee joints were treated with antibiotic and bandaged for the oACh animals, and a plaster cast was additionally applied on the oBMSC animals. Plaster casts were introduced in the oBMSC group, in an effort to further reduce leaning on the operated joint, which was observed with the first group of animals (oACh group). Both groups of sheep recovered in slings for 2 weeks in an effort to limit the weight on the repaired joint, and then were permitted to roam freely for the remaining 6 weeks. A total of 8 weeks after the procedure, sheep were euthanised and joints recovered. The defect regions were trimmed with a saw and fixed in 4% PFA for 72 hours. The defects were decalcified using a KOS rapid decalcification machine (Milestone). The tissues were dehydrated, paraffin embedded, and sectioned at 5 μm. The tissues were stained with hematoxylin and eosin (H&E), as well as Alcian blue and Nuclear Fast Red.

### Statistical analysis

Statistical significance of quantitative data was performed using ANOVA in GraphPad Prism. Statistical significance was set at *P*<0.05. For GAG, DNA, and qPCR analysis, each culture was performed for 4 replicate wells (n=4). The number of unique cell donors or animals were as indicated in the Methods and Results.

## Results

### oBMSC characterisation

Isolated oBMSC appeared spindle-shaped upon expansion (Figure 1A), resembling BMSC from other species such as hBMSC. Flow cytometry analysis is summarised in Figure 1B for oBMSC populations derived from three unique donors, one sheep bone marrow mononuclear cell (MNC) donor population, and one hBMSC donor population (histograms are shown in Supplementary Figures 1-5). Unlike hBMSC, oBMSC showed only a small population of cells positive for CD73 (10.2%±12.7%), negative for CD90 and CD105, and contained a small population positive for CD34 (13.1%±3.7%). Like hBMSC, oBMSC were 100% positive for CD44, mostly positive for CD146 (69%±8.4%), and a small population was positive for CD271 (17.0%±4.3%). The low frequency of CD45^+^ cells (3.1%±1.1%) in oBMSC cultures, along with data that 76% of sheep MNCs stained positively for CD45, suggests that CD45 cells are depleted during the expansion process. Given the successful depletion of CD45 cells during oBMSC enrichment and expansion, the small population of CD34^+^ cells (13.1%±3.7%) likely reflects non-specific binding with this antibody. While the antibody panel had limited cross-reactivity with oBMSC, the relative enrichment of CD146^+^ cells [20] and depletion of CD45^+^ cells [21] was encouraging for future use in characterisation.

**Figure 1.**
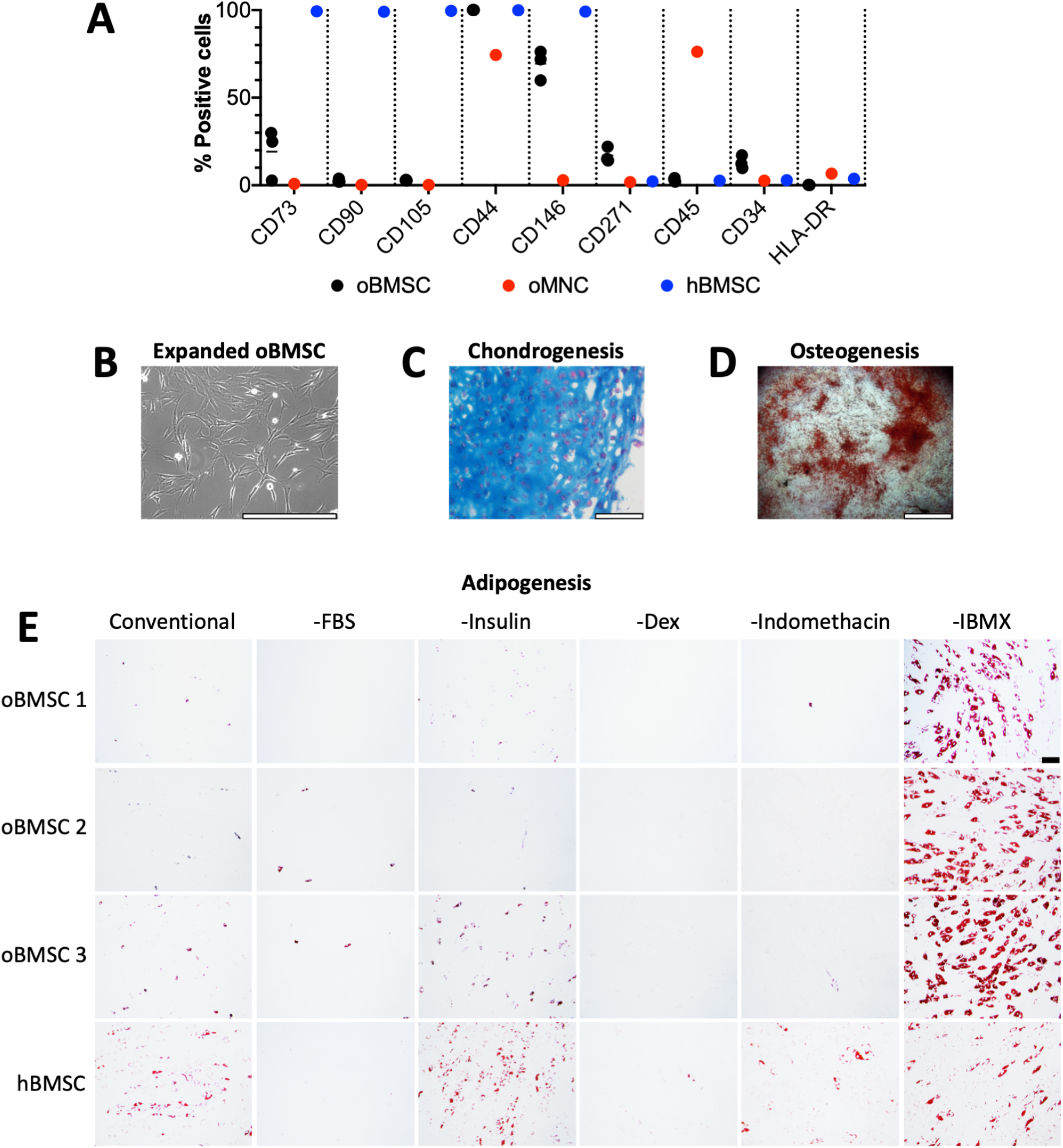
oBMSC characterisation. **A)** Bright field image of oBMSC during expansion. **B)** Flow cytometry analysis of three unique oBMSC donors, with ovine MNC (oMNC) and hBMSC as control populations. Flow cytometry histograms are shown in Supplementary Figures 1-5. **C)** Chondrogenic induction shown by Alcian blue staining of matrix GAG. **D)** Osteogenic induction of oBMSC showing mineralised nodules stained with Alizarin Red S. **E)** Adipogenic induction of 3 unique oBMSC donors and an hBMSC donor, showing that conventional induction media does not result in adipogenesis of oBMSC, unless IBMX is removed. Lipid vacuoles are stained red with Oil Red O. Scale bar = 400 μm in A) and C), 2 mm in D), and 100 μm in E).

The multipotency of oBMSC was evaluated with tri-lineage differentiation in vitro, following protocols conventionally used for hBMSC. Alcian blue staining of oBMSC *macro*-pellets cultured in chondrogenic induction medium demonstrated deposition of cartilage-like extracellular matrix (ECM; Figure 1C). Alizarin Red S staining confirmed calcium nodules in osteogenically induced cultures (Figure 1D). Following adipogenic induction using conventional induction medium, we noticed that adipogenesis was poor and the cells appeared unhealthy and detached from the well plates. To test whether there was a component in the conventional adipogenic medium that was toxic to oBMSC, we set up an experiment in which we removed one component at a time from the conventional adipogenic medium. We also included an hBMSC population in this test as a control. We found that removal of IBMX resulted in healthy oBMSC cultures that underwent adipogenesis, forming ample lipid vacuoles, demonstrated by Oil Red O staining (Figure 1E). Removal of IBMX from hBMSC adipogenic induction cultures did not have the same effect.

### *Growth of* macro*-pellets and* micro*-pellets in the* microwell-mesh *system*

To characterise the chondrogenic potential of oBMSC, we assessed chondrogenesis in both our custom *Microwell-mesh* culture platform (5×10^3^ cells/*micro*-pellet, Figure 2B), as well as traditional *macro-*pellet culture (2×10^5^ cells/*macro*-pellet, Figure 2A). Over 14 days of chondrogenic induction culture, oBMSC *micro*-pellets and *macro-*pellets in 2%, 5%, and 20% O_2_ atmospheres showed an increase in size (Figure 2C). *Micro*-pellets cultured at 2% O_2_ were larger than *micro*-pellets cultured at 5% O_2_ or 20% O_2_, and this was consistent across 3 unique oBMSC donors (see Supplementary Figure 6 for two additional donors). Some variability was observed across different oBMSC donors, particularly in *macro*-pellet cultures (compare Figure 3D with Supplementary Figures 6A and 6C). However, this is not surprising as donor variability has been well-documented with hBMSC [22].

**Figure 2.**
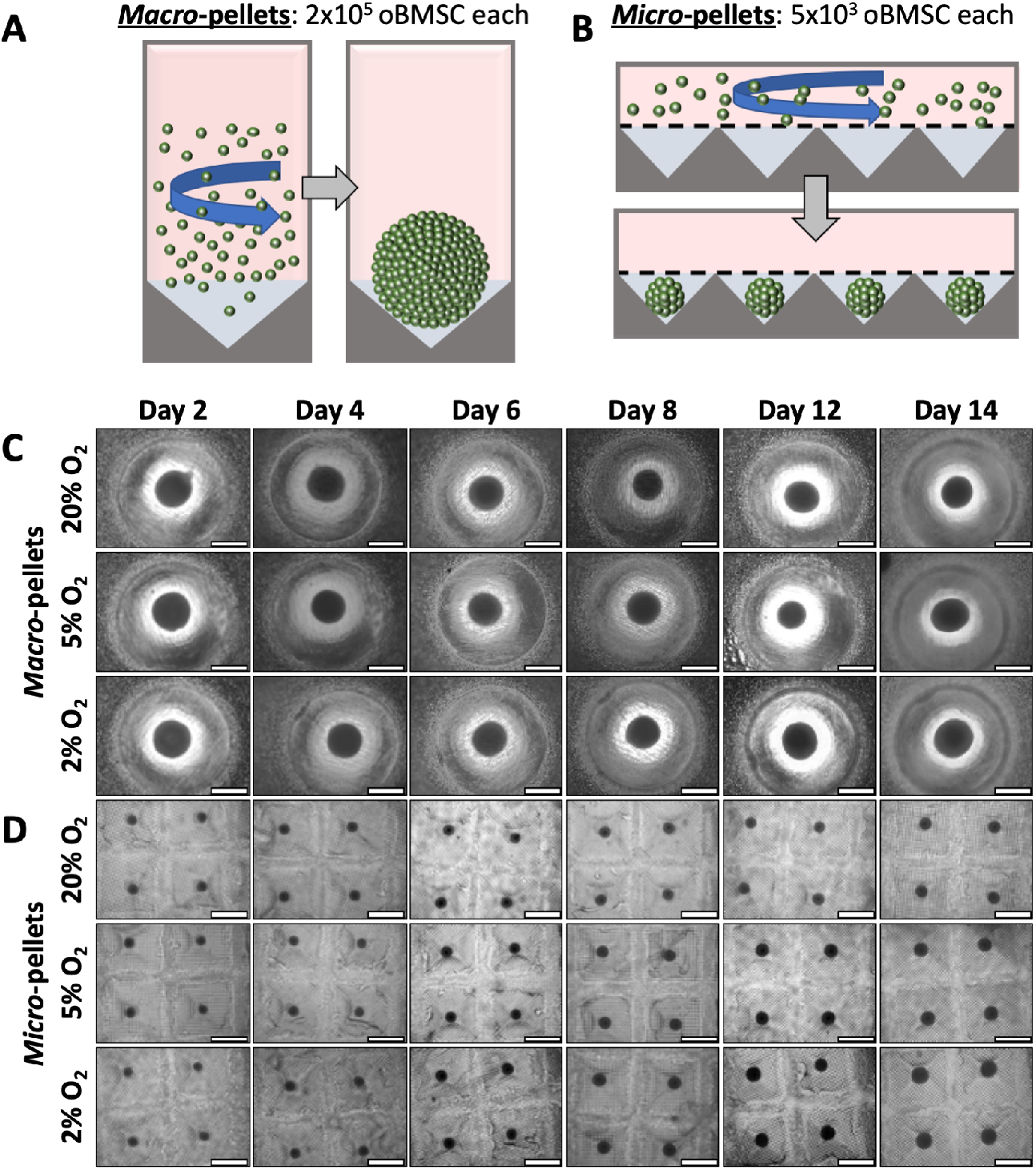
Schematic of *macro*-pellet and *micro*-pellet assembly. **A)***Macro*-pellets were assembled by centrifuging 2×10^5^ oBMSC in V-bottom deep well plates. **B)***Micro*-pellets were assembled by centrifuging 5×10^3^ oBMSC per *micro*well, in the *Microwell-mesh* (A and B adapted from [7]). Microscopic images over a 14-day culture period of **A)***macro*-pellets cultured in deep-well plates and **B)***micro*-pellets cultured in the *Microwell-mesh* platform. Scale bar = 1 mm. Images for 2 additional oBMSC donors are shown in Supplementary Figure 6.

**Figure 3.**
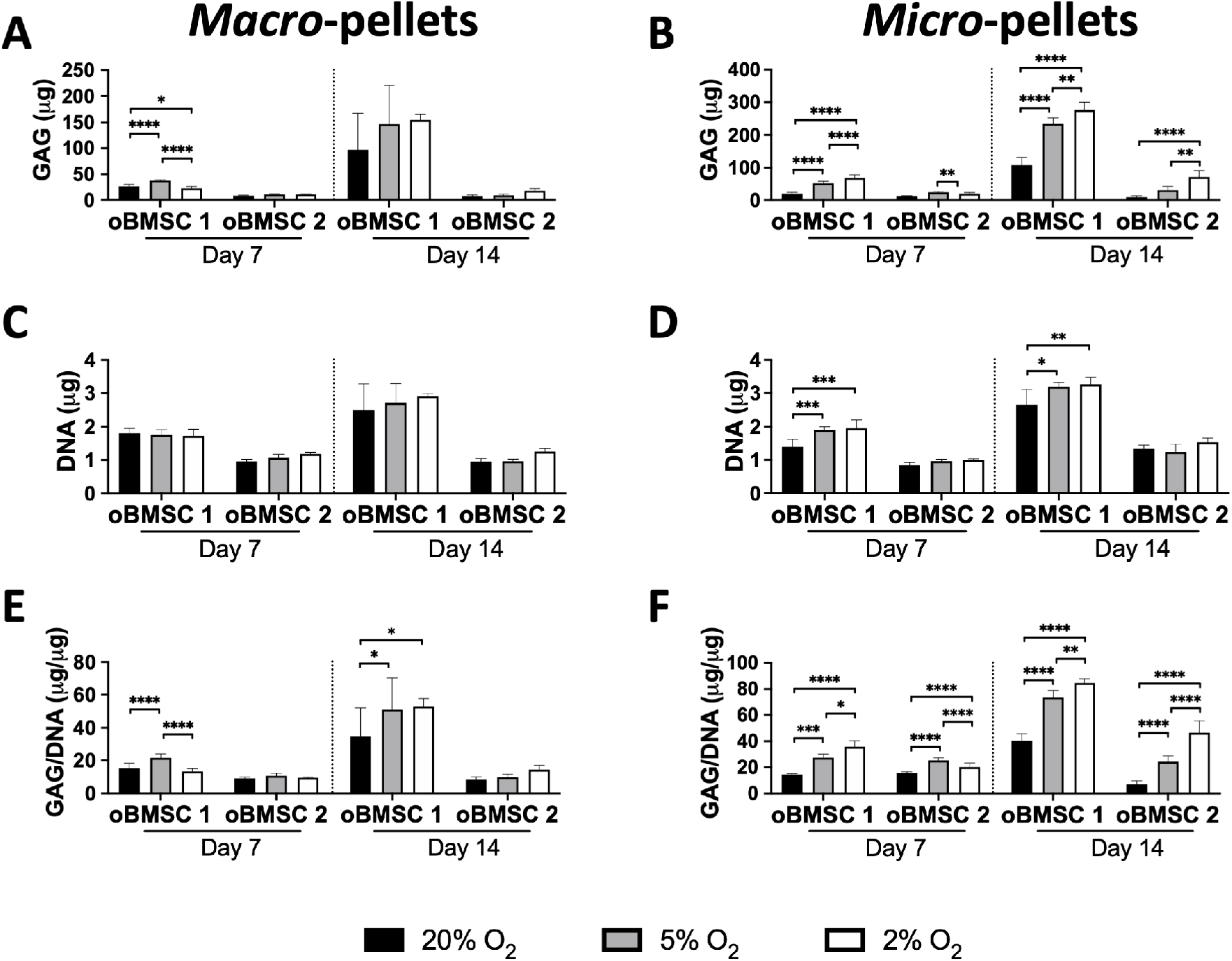
Quantification of GAG and DNA in *macro*-pellets and *micro*-pellets on Day 7 and Day 14 for oBMSC 1 and oBMSC 2. Quantities of GAG in **A)***macro*-pellets and **B)***micro*-pellets. Quantities of DNA in **C)***macro*-pellets and **D)***micro*-pellets. GAG normalised to DNA in **E)***macro*-pellets and **F)***micro*-pellets. (\**P*<0.05, \*\**P*<0.01, \*\*\**P*<0.001, and \*\*\*\**P*<0.0001).

### *Low oxygen increases GAG production in* micro*-pellets*

GAG production increased from Day 7 to Day 14 for both *macro*-pellets and *micro*-pellets, for both of the two unique oBMSC donors analysed, oBMSC 1 and oBMSC 2 (Figure 3A and B). In *macro*-pellet cultures, the amount of GAG and DNA trended toward a higher level at lower O_2_ concentrations (2% and 5%), but this was not consistently statistically significant due to a large amount of variability between individual *macro*-pellets (Figure 3A, C, and E). In *micro*-pellet cultures, the amount of GAG was greater at lower O_2_ concentrations (2% and 5%) and this was statistically significant for oBMSC 1 on Day 7 and Day 14, and for oBMSC 2 on Day 14 (Figure 3B). The amount of DNA in *micro*-pellets was higher for oBMSC 1 at lower O_2_ concentrations (2% and 5%), compared with 20% O_2_, but the magnitude of differences between O_2_ concentrations was generally small (Figure 3D). When GAG was normalised to DNA for *micro*-pellet cultures, the amount of GAG/DNA was greater at lower O_2_ concentrations (2% and 5%), compared with 20% O_2_, and this was statistically significant for both oBMSC donors at both time points (Figure 3F). There was substantial variability observed between the two oBMSC donors, with oBMSC 1 producing substantially more GAG than oBMSC 2 when assessed at both Day 7 and Day 14.

As previously observed for hBMSC [3, 7], GAG/DNA production in *micro*-pellet cultures was typically greater than in *macro*-pellet cultures. At 5% O_2_, GAG/DNA was 1.44±0.11-fold greater (*P*=0.0039) in oBMSC 1 *micro*-pellet cultures than *macro*-pellet cultures on Day 14; and for oBMSC 2, GAG/DNA was 2.27±0.42-fold (*P*<0.0001) greater in *micro*-pellet cultures than *macro*-pellet cultures on Day 7. At 2% O_2_, GAG/DNA was greater in *micro*-pellet cultures than in *macro*-pellet cultures for both donors, and on both Day 7 and 14. For oBMSC 1 at 2% O_2_, GAG/DNA was 2.68±0.34-fold (*P*<0.0001) and 1.60±0.06-fold (*P*<0.0001) greater in *micro*-pellet cultures compared with *macro*-pellet cultures on Day 7 and 14, respectively. For oBMSC 2 at 2% O_2_, GAG/DNA was 4.90±0.96-fold (*P*<0.0001) greater on Day 7 and 3.22±0.63-fold (*P*<0.0001) greater on Day 14 in *micro*-pellet cultures compared with *macro*-pellet cultures.

### *ECM characterisation in* macro*-pellets and* micro*-pellets*

We assessed the production of cartilage-like ECM with 3 unique oBMSC donors in *macro*-pellets and *micro*-pellets cultured in 2%, 5%, and 20% O_2_ atmospheres, on Day 7 and Day 14. Alcian blue staining revealed increased GAG matrix accumulation in both *macro*-pellets and *micro*-pellets cultured at 2% O_2_ compared with 20% O_2_ (Figure 4A and B). On Day 7, *macro*-pellets contained cell-dense cores (red stained nuclei) with little cartilage-like matrix at all oxygen levels (Figure 4A). By Day 14, *macro*-pellets exhibited more uniform GAG matrix throughout their diameter for oBMSC 1 at all oxygen concentrations, and for oBMSC 2 and oBMSC 3 at 2% O_2_. Similarly, *micro*-pellets cultured at 2% O_2_ stained more uniformly with Alcian blue throughout their diameter by Day 14, compared with 5% and 20% O_2_ (Figure 4B). Consistent with GAG quantification (Figure 3), variability was also evident in Alcian blue GAG staining between unique oBMSC donors.

**Figure 4.**
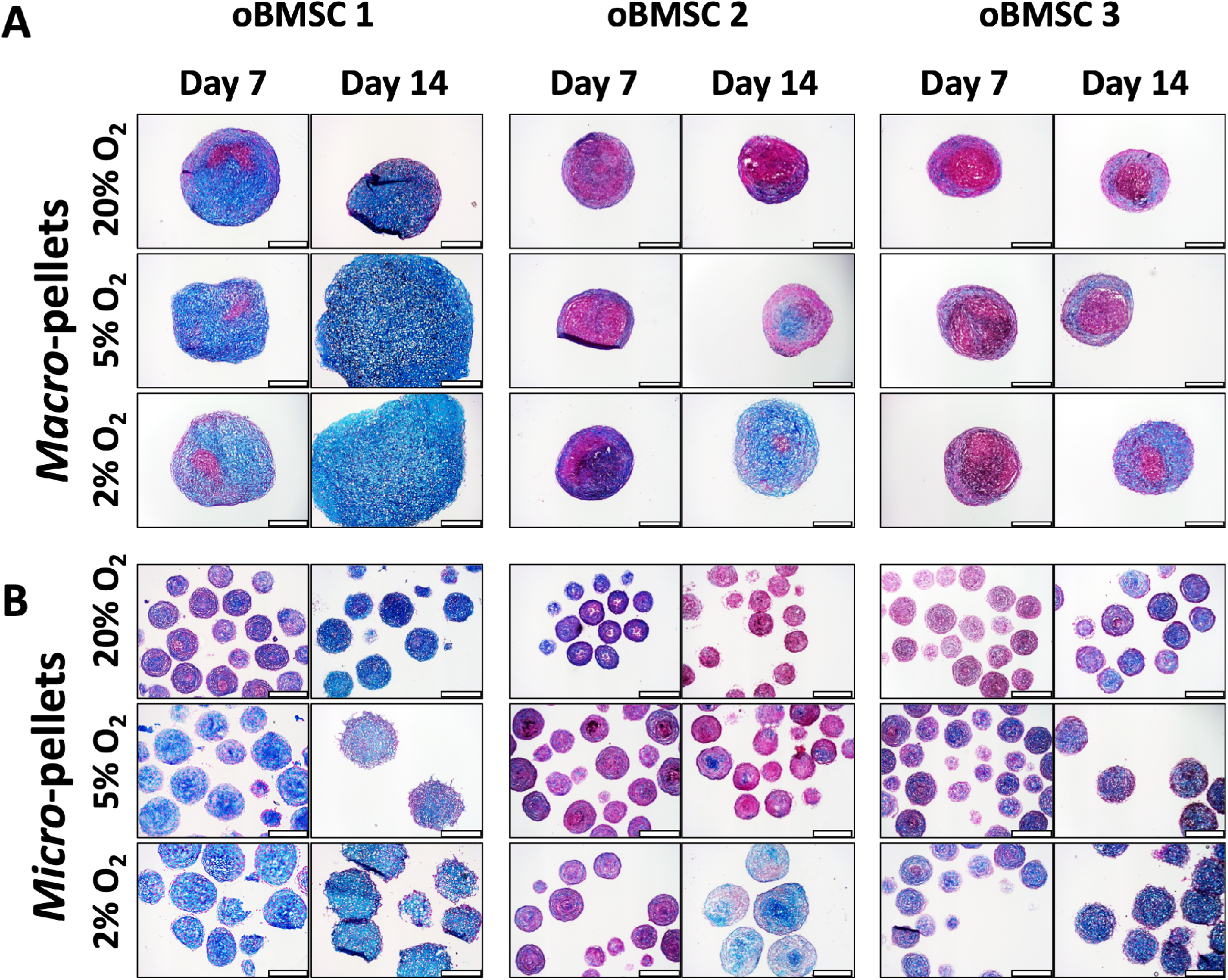
Alcian blue staining for GAG in **A)***macro*-pellet and **B)***micro*-pellet sections. Nuclei are stained red. Scale bar = 400 μm.

We also performed immunohistochemistry staining for cartilage-associated ECM molecule, type II collagen (Supplementary Figure 7); bone-associated molecule, type I collagen (Supplementary Figure 8); and hypertrophy-associated molecule, type X collagen (Supplementary Figure 9). While we observed some staining for all of these collagen molecules, we did not observe a staining pattern that correlated with differing oxygen concentrations.

### In vivo *cartilage defects in sheep*

We evaluated the potential of oBMSC *micro-*pellets to repair cartilage defects in sheep. *Micro*-pellets induced for 10 days were suspended in fibrin glue and anchored to a collagen Celgro scaffold/membrane that was used to keep the tissues fixed in osteochondral defects in sheep (see schematic in Figure 5). Three defects (6 mm in diameter and 1.5 mm deep) were made in the trochlear groove of one hind leg for each sheep and filled with repair tissues. Two sheep were implanted with *micro*-pellets made from induced oBMSC, and two sheep were implanted with *micro*-pellets made from induced oACh, as controls. Sheep were euthanised after a total of 8 weeks from the day of receiving an implant, and defect repair was assessed by histology.

**Figure 5.**
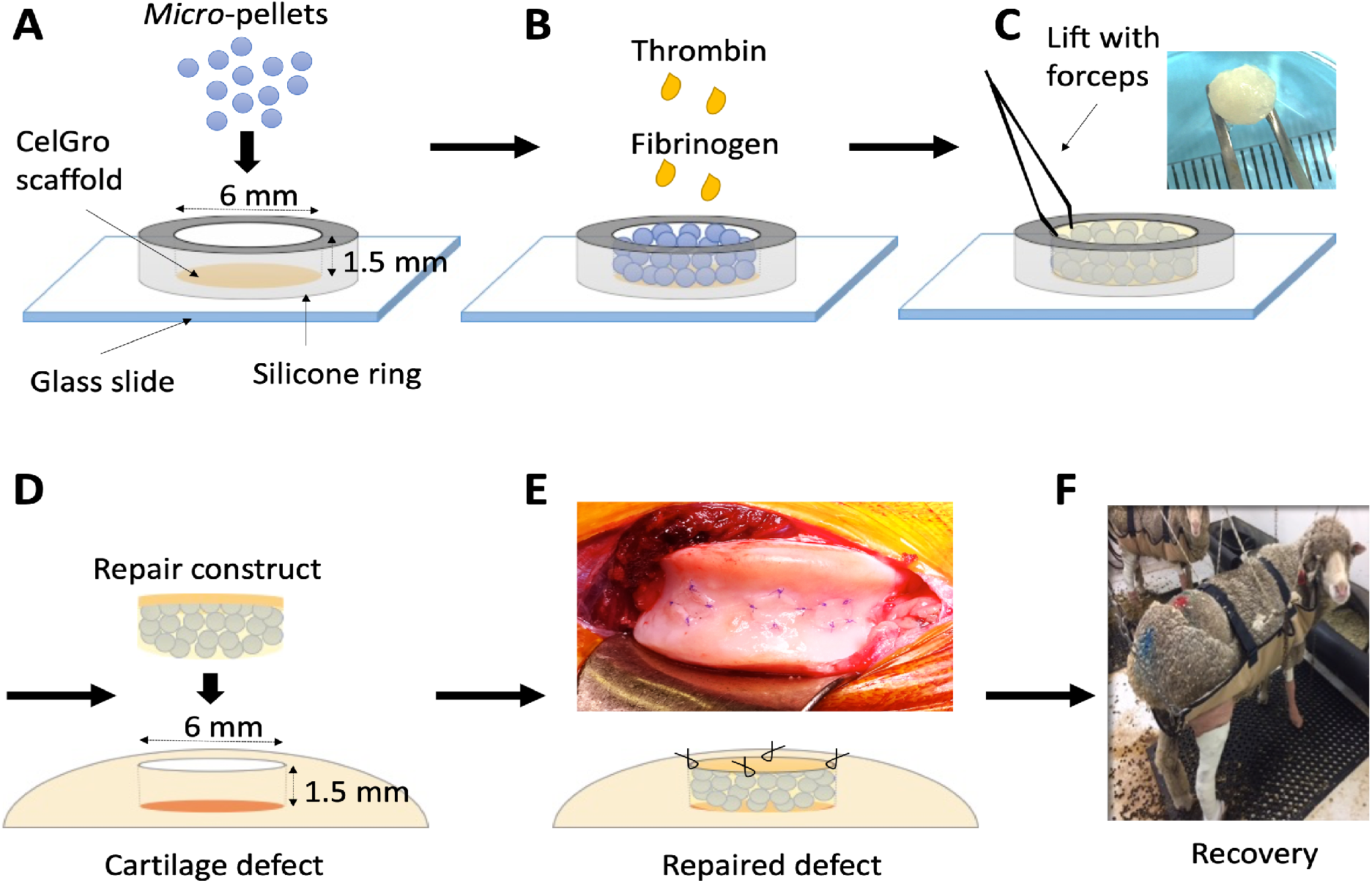
Schematic of *in vivo* evaluation of *micro*-pellets. **A)** Ten-day chondrogenically induced oBMSC or oACh *micro*-pellets were layered onto a Celgro scaffold in a custom-made cylindric chamber (6 mm in diameter and 1.5 mm high). **B)** Two drops of fibrinogen were added to the *micro*-pellets, followed by two drops of thrombin to adhere the *micro*-pellets to each other and to the Celgro scaffold. **C)** The repair construct was lifted with forceps. **D)** The repair construct was placed into a cartilage defect that was of a similar dimension, with the *micro*-pellet layer facing down into the subchondral layer and the Celgro scaffold on top. **E)** Four sutures were used to fix the Celgro scaffold to the adjacent cartilage tissue. **F)** Sheep were placed in slings for 2 weeks to recover and then were allowed to roam freely for the remaining 6 weeks.

**Figure 6.**
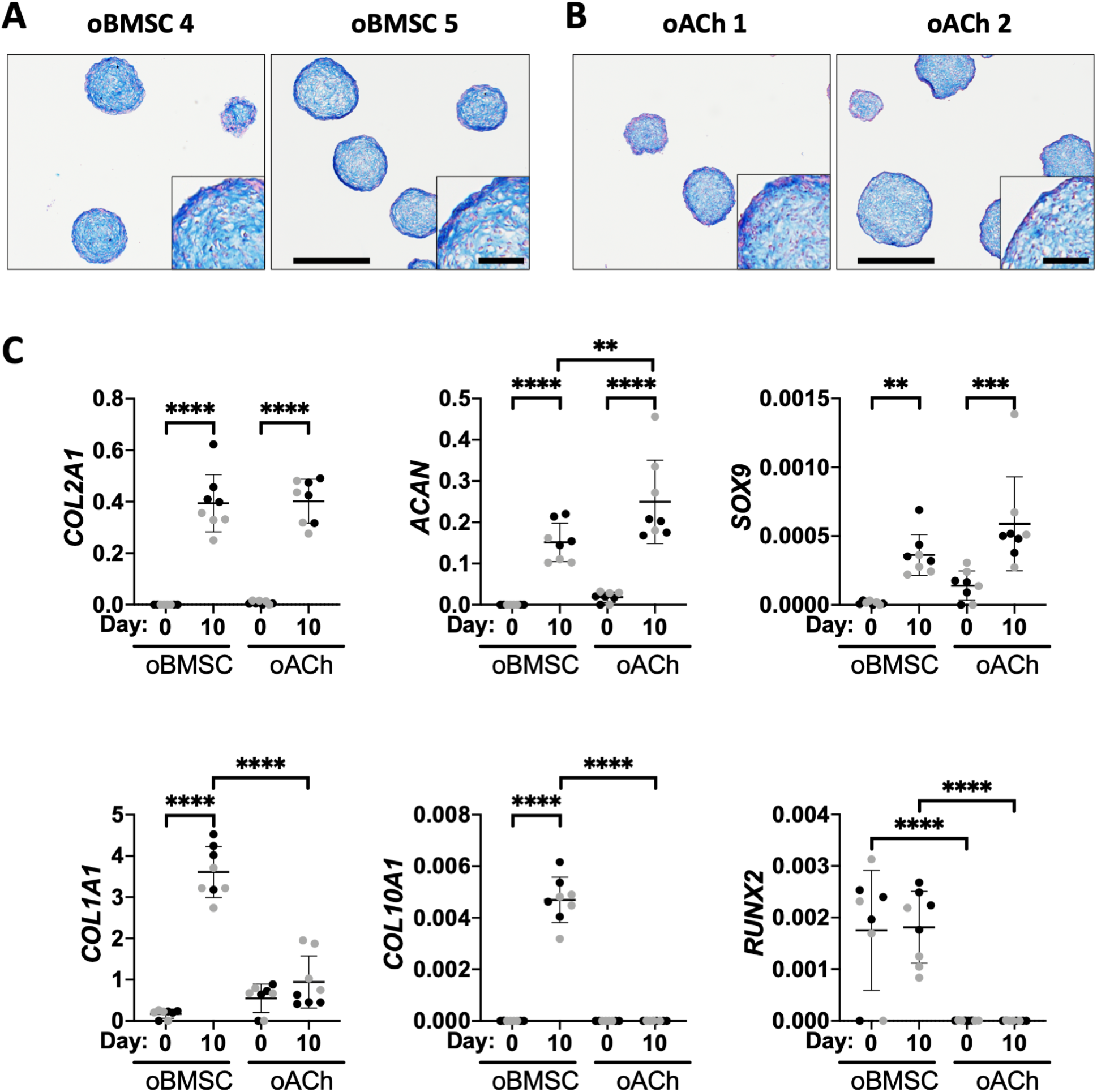
Characterisation of induced *micro*-pellets used for sheep studies. **A)** oBMSC and **B)** oACh *micro*-pellet sections stained with Alcian blue for GAG following 10 days of chondrogenic induction (scale bar = 500 μm and inset scale bar = 100 μm). **C)** qPCR analysis of cartilage-like tissues formed from oBMSC and oACh after expansion (Day 0) and 10 days after chondrogenic induction (Day 10). Relative gene expression was assessed for chondrogenic genes *COL2A1*, *ACAN*, and *SOX9*, and for hypertrophic genes *COL1A1*, *COL10A1*, and *RUNX2*. Gene expression levels were normalised to *GAPDH* and represent ΔCt values. The two unique donors are distinguished by black versus grey symbols. The mean is represented by the horizontal line (\**P*<0.05, \*\**P*<0.01, \*\*\**P*<0.001, and \*\*\*\**P*<0.0001).

### *Histological and qPCR assessment of oBMSC and oACh* micro*-pellets pre-implant*

We assessed the *micro*-pellets implanted in sheep histologically and for relative gene expression (Figure 6). Ten-day, chondrogenically induced *micro*-pellets derived from both oBMSC (Figure 6A) and oACh (Figure 6B) donors stained for GAG with Alcian blue, demonstrating the development of cartilage-like tissue. Histologically, the oBMSC- and oACh-derived *micro*-pellets were visually indistinguishable from each other. qPCR analysis was performed to quantify the relative gene expression (Figure 6C) for cartilage-associated genes (*COL2A1*, *ACAN*, and *SOX9*) and for bone or hypertrophy-associated genes (*COL1A1, COL10A1*, and *RUNX2*) in both oBMSC- and oACh-derived *micro*-pellets, at Day 0, before induction, and on Day 10 of induction. For both oBMSC- and oACh-derived *micro*-pellets, the expression of *COL2A1*, *ACAN*, and *SOX9* were statistically significantly increased on Day 10 following chondrogenic induction, relative to Day 0. For oBMSC-derived tissues, *COL2A1*, *ACAN*, and *SOX9* increased by 1.45×10^6^ ± 3.82×10^5^-fold, 2.56×10^4^ ± 7.38×10^3^-fold, and 61.3 ± 23.6-fold, respectively, on Day 10 relative to Day 0. For oACh-derived tissues, *COL2A1*, *ACAN*, and *SOX9* increased by 53.8 ± 10.7-fold, 13.5 ± 5.11-fold, and 4.23 ± 2.29-fold, respectively, on Day 10 relative to Day 0. *ACAN* expression was 1.65±0.624 higher in oACh *micro*-pellets following induction (Day 10) relative to oBMSC *micro*-pellets (Day 10), whereas *COL2A1* and *SOX9* were statistically similar between oBMSC and oACh at both time points. For oBMSC-derived tissues, the expression of *COL1A1* (21.3 ± 3.41-fold) and *COL10A1* (1.61×10^3^ ± 283-fold) were increased following chondrogenic induction, while *RUNX2* expression was not changed following induction. Following induction (Day 10), *COL1A1* (8.97±1.44-fold) and *COL10A1* (1.10×10^3^ ± 193-fold) were higher in oBMSC-derived *micro*-pellets relative to oACh-derived *micro*-pellets. *RUNX2* expression was higher in oBMSC-derived *micro-*pellets compared with oACh *micro*-pellets, both on Day 0 (345 ± 213-fold) and on Day 10 (708 ± 254-fold).

### *Histological assessment of oBMSC and oACh* micro*-pellets post-implant*

Following 8 weeks of incubation *in vivo* in sheep, joints were harvested, and defect repair was characterised histologically (Figures 7 and 8). None of the oBMSC or oACh *micro*-pellet groups showed complete cartilage fill or seamless integration with adjacent tissue. Figure 7 shows three representative defects that were filled with oBMSC-derived *micro*-pellets and stained with H&E and Alcian blue. H&E staining showed that most of the defects were filled with fibrous tissue, vascular tissue, and blood cells (Figure 7A-C). There was faint Alcian blue staining for GAG in some of the defects (see Figure 7B, lower left panel), suggesting either residual cartilage from the implanted *micro*-pellets, or that cartilage repair was occurring very slowly. This could also have been from infiltration of chondroprogenitors from adjacent tissue rather than oBMSC. Supplementary Figure 10 shows histology of a scaffold-only control, where the defect was filled with fibrin glue and the Celgro scaffold, but no cells. Of the 5 defects filled with oBMSC *micro*-pellets, we could only find one defect that still had some recognisable *micro*-pellets, which are shown Figure 7C. Based on the morphology of these oBMSC *micro*-pellets, it appeared as though they may have been in the process of dedifferentiating or being absorbed by surrounding tissue.

**Figure 7.**
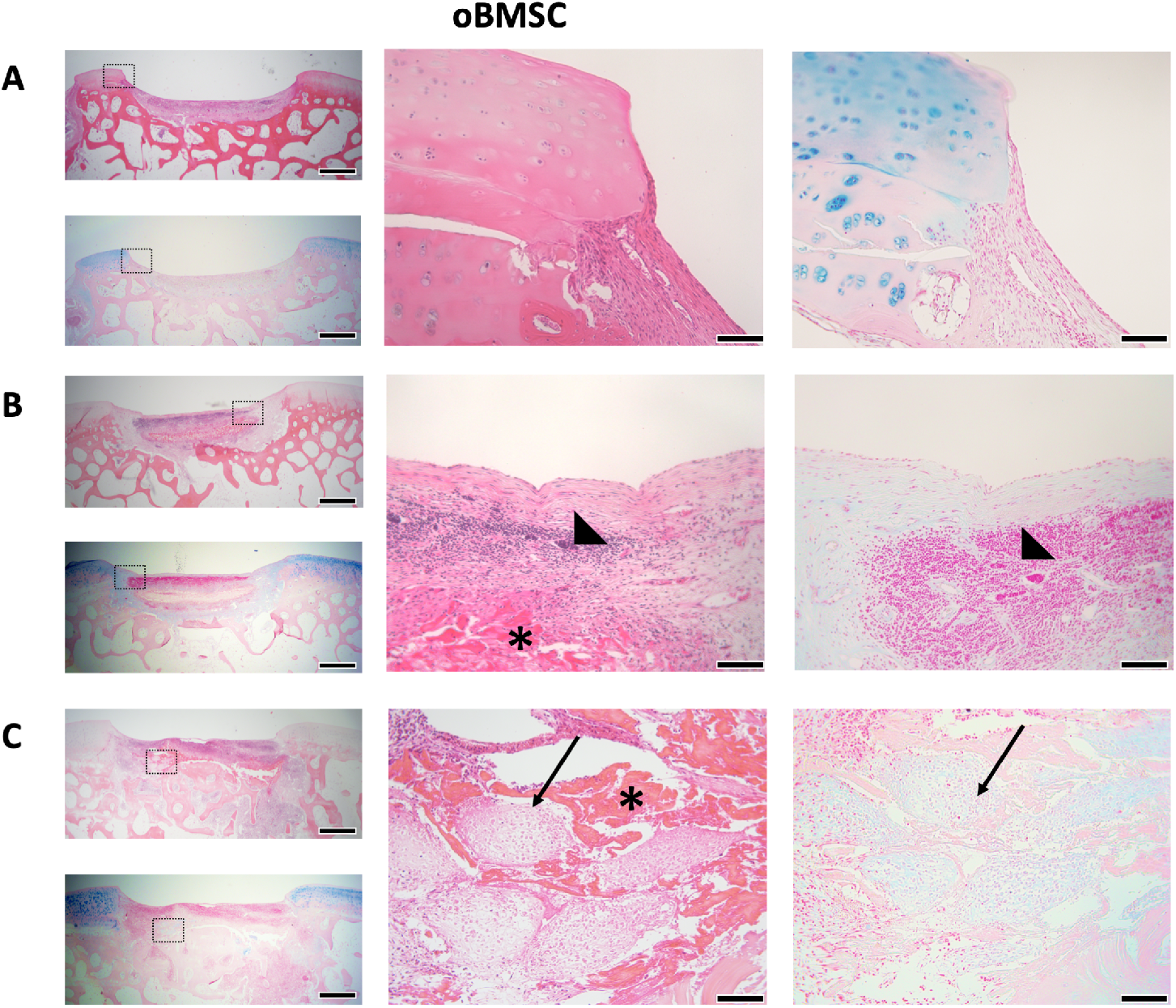
Representative H&E and Alcian blue stained images of *in vivo* oBMSC *micro*-pellet repair tissues after 8 weeks. Repair tissue was largely composed of **A)** fibrous tissue and **B)** fibrous tissue with significant vascular and blood cell infiltration (triangle). **C)** In one of 6 defects, a few *micro*-pellets could be identified; the boxed regions are enlarged to the right, with arrows pointing to *micro*-pellets. The scaffold was largely degraded, but could be seen in some defects (see asterisks in B and C). Boxes with dashed lines in images on the left are enlarged in the middle (H&E) and right (Alcian blue) images. Scale bars in left panels = 1 mm. Scale bars in middle and right panels = 100 μm.

**Figure 8.**
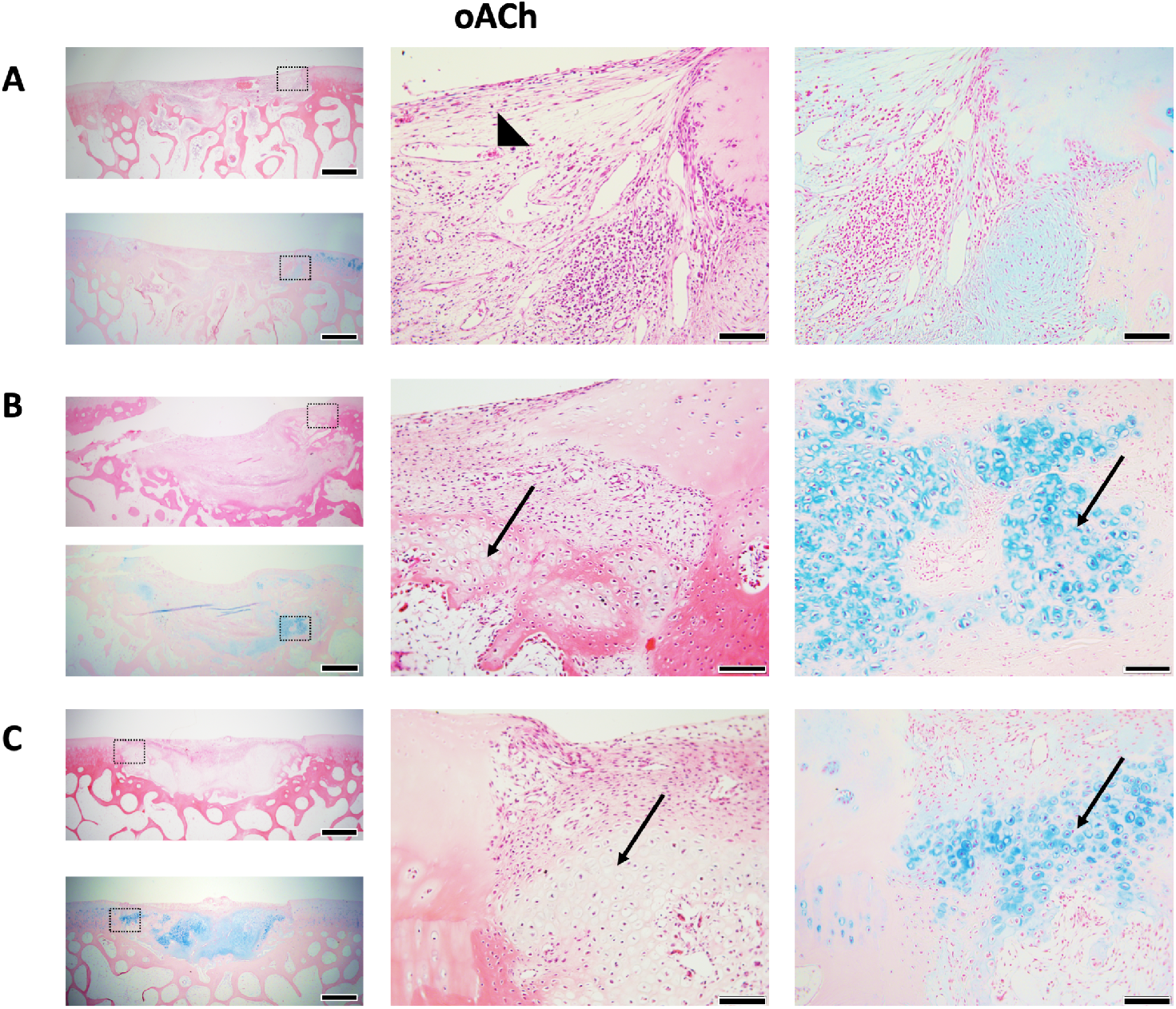
Representative H&E and Alcian blue–stained images of *in vivo* oACh *micro*-pellet repair tissues after 8 weeks. Repair tissue was largely composed of **A)** fibrous tissue with some vascular tissue (triangle), with **B)** one defect showing modest regions of GAG-rich repair cartilage (arrows), and **C)** another defect showing more substantial regions of GAG-rich repair cartilage (arrows). Boxes with dashed lines in images on the left are enlarged in the middle (H&E) and right (Alcian blue) images. Scale bars in left panels = 1 mm. Scale bars in middle and right panels = 100 μm.

Figure 8 shows three representative defects that were filled with oACh-derived *micro*-pellets and stained with H&E and Alcian blue. H&E staining showed that most of the defect was filled with fibrous tissue, but unlike defects filled with oBMSC *micro*-pellets, they generally appeared to have attracted fewer blood lineage cells and less vascular tissue. Of the 5 defects filled with oACh *micro*-pellets, we could only find two defects that still had substantial cartilage-like tissue fill, which are shown in Figure 8B and 8C. There was still a significant amount of fibrous tissue observed in these defects, however, they showed the most promising result in terms of the amount of Alcian blue– stained GAG matrix and morphology of the cells, which appeared to form lacunae. Supplementary Figure 10 shows histology from an empty Scaffold-only control defect site. While oACh-derived *micro*-pellets yielded more promising histology, macroscopic imaging of the joint revealed that defects remained visible, with substantial repair visible only in a single defect (Supplementary Figure 11).

## Discussion

In this study we characterised oBMSC expansion cultures *in vitro*, used a *micro*-pellet model to identify the optimal oxygen atmosphere for oBMSC chondrogenesis, and then compared the capacity of *micro*-pellets assembled from either oBMSC or oACh to repair osteochondral defects in adult sheep.

oBMSC were successfully isolated using plastic adhesion and cells exhibited a spindle-shaped morphology like hBMSC. Expanded oBMSC were characterised using a flow cytometry antibody panel commonly used to characterise hBMSC [23, 24], including CD73, CD90, CD105, CD44, CD146, CD271, CD45, CD34, and HLA-DR. Like our control hBMSC population, expanded oBMSC were essentially negative for haematopoietic markers (CD45 and CD34), strongly positive (~100%) for CD44, contained a large population (~70%) of CD146^+^ cells, and had a very small population (<3%) of CD271^+^ cells. CD146 is viewed as one of the defining markers of hBMSC, and is associated with hBMSC capacity to form ectopic marrows and support haematopoietic stem cells [25, 26]. CD271 has been reported to be expressed on the surface of hBMSC [27] and oBMSC [28]. However, CD271^+^ cell number in hBMSC cultures declines rapidly in response to medium serum supplementation [29], and our hBMSC control population and oBMSC cultures also had few CD271^+^ cells. We did not observe positive staining of oBMSC using anti-human CD90 or CD105 antibodies (<3%), and only two oBMSC donors stained weakly for CD73 (<30%), while one donor was essentially negative for CD73 (<3%). A previous study also reported poor cross-reactivity of oBMSC surface antigens and anti-human CD73, CD90, and CD105 antibodies [30]. As expected, oBMSC were negative for the human specific marker, HLA-DR, which served as a negative control. Other markers that have been positively detected on oBMSC, include CD166 and CD29 [31]. While robust correlations between BMSC surface antigens and biological function are lacking, these markers remain useful for identifying and comparing stromal cells from different tissues and species.

Using conventional tri-lineage induction media, oBMSC underwent osteogenic and chondrogenic induction successfully, while initially, adipogenic induction failed. Using conventional adipogenic induction medium, we observed substantial cell death. By removing one component of the induction medium at a time, we identified that IBMX was toxic to oBMSC. Exclusion of IBMX from the conventional adipogenic induction medium resulted in robust adipogenesis of oBMSC. IBMX is a competitive non-selective phosphodiesterase inhibitor [32], commonly included in BMSC adipogenic induction medium [33]. While other studies do not specifically report the toxicity of IBMX during oBMSC adipogenesis, a previous publication showed an image of a single cell from hBMSC and oBMSC cultures after 14 days of induction in IBMX-supplemented medium [31, 34]. In these images the adipogenic induced hBMSC appeared healthy and contained many lipid vacuoles, while the oBMSC appeared relatively small with few lipid vacuoles [34]. Another study induced oBMSC in medium supplemented with IBMX, and displayed cells in a relatively sparse monolayer, with few lipid vacuoles [35], potentially indicating some level of toxicity. These outcomes are similar to our own observations where only a few small oBMSC remained in adipogenic cultures supplemented with IBMX. For effective oBMSC adipogenic induction, exclusion of IBMX from induction cultures appears to be necessary. Why oBMSC respond differently to IBMX during adipogenic induction, compared with hBMSC, is not immediately obvious. This toxicity, however, identifies an important difference in the biology of oBMSC and hBMSC, and highlights the importance of characterising and comparing cells of different species independently.

We characterised the capacity of oBMSC to form cartilage-like tissue in *micro*-pellet and traditional *macro*-pellet cultures, in 2%, 5%, or 20% O_2_ atmospheres. A previous study that included both an oBMSC pellet model and mathematical modelling concluded that optimal oBMSC chondrogenesis occurs at 10-11% O_2_ [36]. Another previous study demonstrated that superior chondrogenesis could be achieved if the oBMSC were expanded in a 5% O_2_ atmosphere, rather than at 20% O_2_ atmosphere [37]. In this previous study, oBMSC expanded in 5% or 20% O_2_ atmospheres were assembled into *macro*-pellets for chondrogenic differentiation [37]. In our study, we expanded all oBMSC in a 2% O_2_ atmosphere, and observed incrementally greater GAG production when oBMSC were assembled into a *micro*-pellet model and cultured at 2% O_2_, compared with 5% or 20% O_2_ atmospheres. GAG/DNA output was greatest in *micro*-pellets cultured in 2% O_2_ atmosphere. In *macro*-pellet cultures, GAG/DNA content trended toward higher levels in lower O_2_ atmospheres, but this was not consistently statistically significant, and greater variability in data was observed in *macro*-pellet cultures. At 2% O_2_, GAG/DNA was significantly greater in *micro*-pellet cultures relative to *macro*-pellet cultures. This pattern of improved chondrogenesis both in *micro*-pellets and at 2% O_2_ was previously observed in hBMSC chondrogenic cultures [3]. For both oBMSC (in this study) and hBMSC (previous studies [3, 7]), more homogeneous tissue was generated using *micro*-pellet cultures, relative to *macro*-pellet cultures. The effect of atmospheric O_2_ concentration was more discernible in homogeneous *micro*-pellet models, demonstrating that, as with hBMSC chondrogenesis, oBMSC chondrogenesis appears to be more efficient at lower O_2_ concentrations.

Next, we sought to evaluate the use of autologous oBMSC *micro*-pellets in an *in vivo* sheep pilot study. Since autologous ACh are already being used in various clinical cartilage repair strategies [38], we included expanded oACh *micro*-pellets for comparison. A 10-day chondrogenic induction culture, rather than 14 days, was selected based on a previous study that indicated that following ~10 days of culture, *micro*-pellets gained significant mass and yet still retained their propensity to amalgamate into a continuous tissue [10]. We characterised oBMSC and oACh *micro*-pellets from 2 unique donors each, using histology and qPCR. By Day 10, both oBMSC and oACh *micro*-pellets stained for GAG with Alcian blue, and both tissues exhibited similar tissue morphology. Conversely, qPCR analysis revealed that hypertrophic gene expression (*COL10A1* and *RUNX2*) was absent in oACh *micro*-pellets, but significantly over-represented in oBMSC *micro*-pellets. Hypertrophic signalling and upregulation of endochondral ossification pathways is a well-characterised problem in hBMSC chondrogenic cultures, and is viewed as a major impediment to BMSC-mediated cartilage repair [7, 39]. It is common to demonstrate hBMSC hypertrophy by implanting tissue ectopically in immune compromised mice [3, 7, 25, 40]. However, it is unclear if the highly vascularised ectopic microenvironment exacerbates the propensity of hBMSC to undergo hypertrophy, and if hypertrophy would be less problematic in the less vascular environment found in a synovial joint. For this reason, there is merit in evaluating the capacity of BMSC to regenerate cartilage in actual cartilage defect models.

To assess the potential of *micro*-pellets to repair osteochondral defects, *micro*-pellets were packed into cylindrical molds, bonded to a Celgro™ collagen membrane with fibrin glue, and then anchored into defect sites with fibrin glue and sutures (see Figure 5). During post-surgical recovery, slings were used to partially reduce weight bearing on the treated sheep joints and plaster casts were introduced to prevent joint mobility. Our ethics protocol did not allow for complete unweighting of the repaired joint. In clinical cartilage repair procedures such as MACI (reviewed here [41]), human patients are advised to adhere to strict progressive weight bearing and passive motion following these procedures. Because the sheep knee joint is high up on the animal’s flank, it is challenging to immobilise this joint with wrapping or a cast. Despite the formation of three 6 mm diameter osteochondral defects in one stifle joint, animals did not appear to be in pain or appear to reduce load on the repaired leg while in the sling or after release from the sling. Full unloading of the treated joint for one week or more could potentially improve engraftment and the stability of repair tissues.

Following 8 weeks of *in vivo* incubation, defects filled with oBMSC *micro*-pellets were not effectively repaired. oBMSC *micro*-pellets were visible in 1 of 5 defect sections, but the lacunae structure and Alcian blue staining suggested that the cartilage-like tissue was either de-differentiating or being absorbed by surrounding tissue. H&E staining suggested that most of the defect fill volume contained fibrous tissue, rather than cartilage-like repair tissue. A layer of tissue was stained faintly with Alcian blue at the base of some defect sites, but it is possible that this tissue was derived from adjacent native tissue, or residual oBMSC *micro*-pellets rather than new cartilage tissue. Repair from adjacent tissue might be consistent with the concept that BMSC-based therapies can upregulate endogenous regenerative processes [42]. However, this repair was modest and our limited controls did not allow for a statistical evaluation of this outcome. In previous studies, we implanted hBMSC *micro*-pellets subcutaneously in NSG mice, and after 8 weeks we observed mineralisation and the formation of marrow-like structures [3]. This is consistent with a wide body of literature that suggests that chondrogenically-induced hBMSC appear to engage intrinsic endochondral ossification programming, triggering vascular penetration, ossification and support of marrow formation [39, 43]. However, in this study we observed no evidence of mineralisation or marrow formation from oBMSC *micro*-pellets implanted into osteochondral defect sites. Another study that implanted oBMSC in TGF-β1-laden gelatine foams in sheep tibial growth plates similarly reported a lack of mineralised tissue, with mostly fibrous tissue present after 5 weeks [44]. It is possible that during the incubation period in our studies, *micro*-pellet mineralisation occurred, but that this tissue was completely re-absorbed by immune cells. Alternatively, it is possible that the microenvironment within these joint defect sites might mitigate mineralisation of tissue, relative to more vascularised microenvironments such as a subcutaneous mouse pouch. While we did not see evidence of mineralisation at 8 weeks, oBMSC *micro*-pellets did not yield promising cartilage-like repair tissue.

In the oACh *micro*-pellet group, evidence of cartilage repair tissue was visible in 2 of 5 histology sections. While there was evidence of continuous cartilage-like tissue formation in these regions, there were also regions of fibrous tissue and vasculature. Where cartilage repair was visible, this repair tissue filled both the articular cartilage defect and the subchondral bone region. Mature lacunae structures and Alcian blue staining were visible in this repair tissue. Longer incubation periods beyond 8 weeks would be required to determine if repair tissue integration and a high-quality cartilage repair tissue might evolve. Many sheep studies are carried out for longer periods (8, 10, 12 weeks [45], 3 months [46], 4-12 months [47]), and we presume that promising aspects of repair with oACh *micro*-pellets would have improved with time. The low success rate of establishing a repair tissue in the oACh group may also be improved by further reducing weight-bearing and motion of the joints during the first few weeks of recovery.

## Conclusion

In summary, cell-mediated cartilage repair strategies in the leading large animal cartilage repair model, the sheep, remain in their infancy. Identifying optimal oBMSC culture conditions and understanding the limitations of oBMSC will likely be critical to advancing the field of cartilage defect repair. Our data demonstrate that **(1)** there is limited cross-reactivity between common hBMSC antibody panels and oBMSC, but that CD44, CD146, and CD45 are useful markers; **(2)** IBMX is toxic during oBMSC adipogenic culture, but IBMX is not required and adipogenic induction can be salvaged by excluding this molecule from the medium; **(3)** oBMSC chondrogenesis can be enhanced by using *micro*-pellet cultures and reduced oxygen atmospheres *in vitro*; **(4)***micro*-pellets formed from oBMSC appear morphologically similar to *micro*-pellets formed from oACh, but have upregulated hypertrophic gene expression *in vitro*; and **(5)** oBMSC *micro*-pellets produce inferior cartilage repair tissue *in vivo*, compared to oACh *micro*-pellets.

## Abbreviations

BMSC: bone marrow stromal cells
hBMSC: human BMSC
oBMSC: ovine BMSC
oACh: ovine articular chondrocytes
IBMX: 3-Isobutyl-1-methylxanthine
HIF: hypoxia-inducible factor
FBS: fetal bovine serum
FGF-1: fibroblast growth factor-1
LG-DMEM: low glucose Dulbecco’s Modified Eagle’s Medium
PenStrep: penicillin/streptomycin
EDTA: ethylenediaminetetraacetic acid
MNC: mononuclear cell
Anti-Anti: Antibiotic-Antimycotic
PDMS: polydimethylsiloxane
PBS: phosphate buffered saline
Dex: dexamethasone
HG-DMEM: high glucose Dulbecco’s Modified Eagle’s Medium
PFA: paraformaldehyde
qPCR: quantitative polymerase chain reaction
BSA: bovine serum albumin
H&E: hematoxylin and eosin
GAG: glycosaminoglycans
DNA: deoxyribonucleic acid
ANOVA: Analysis of variance

## Ethics approval and consent to participate

All animal procedures were approved by the Queensland University of Technology Animal Ethics Committee. Human bone marrow was collected from informed consenting healthy volunteer donors. Donation and use of cells were approved by the Mater Hospital Human Research Ethics Committee, and the Queensland University of Technology Human Research Ethics Committee.

## Consent for publication

Not applicable

## Availability of Data and Materials

Additional information or data is available through the Senior Author.

## Author contributions

KF, EM, PGR, SG, RWC, SS, TJK and MRD designed research, analysed data, and wrote the paper; EM, KF, SS and MRD performed research.

## Acknowledgments

The authors thank the core facilities at the TRI, including the Flow Cytometry Core, and the Microscopy Core. The authors thank the surgical team at the Medical Engineering Research Facility (MERF, QUT) for assistance in organising and executing this study. The authors would like to thank Professor MingHao Zheng for technical advice, and Orthocell Ltd (Perth Australia) for donating the Celgro scaffolds used in these studies.

