## Supplementary Material for "Characterisation of ovine bone marrow-derived stromal cells (oBMSC) and evaluation of chondrogenically induced *micro*-pellets for cartilage tissue repair *in vivo*" (from another version of this preprint: version 1, 2020-05-16)

### Supplementary Data

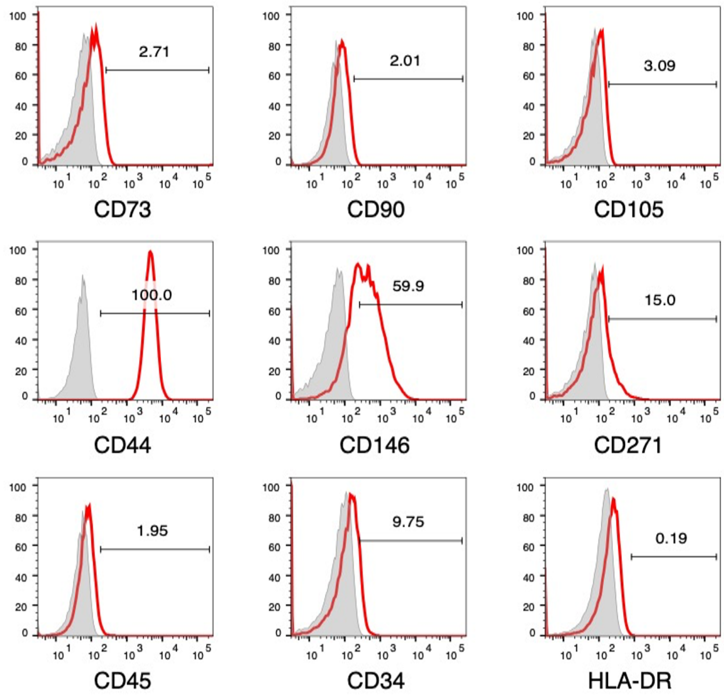


**Supplementary Figure 1.** Flow cytometry analysis of cell surface markers for oBMSC 1.

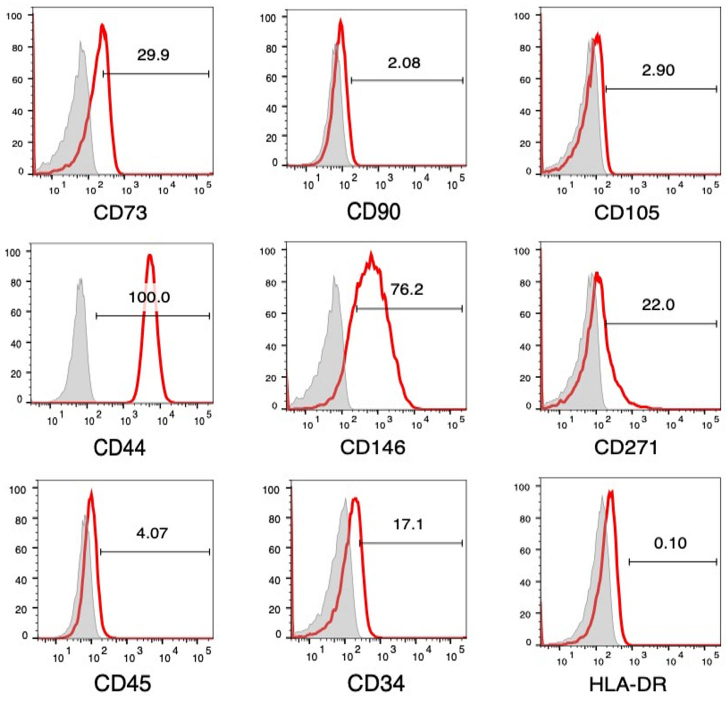


**Supplementary Figure 2.** Flow cytometry analysis of cell surface markers for oBMSC 2.

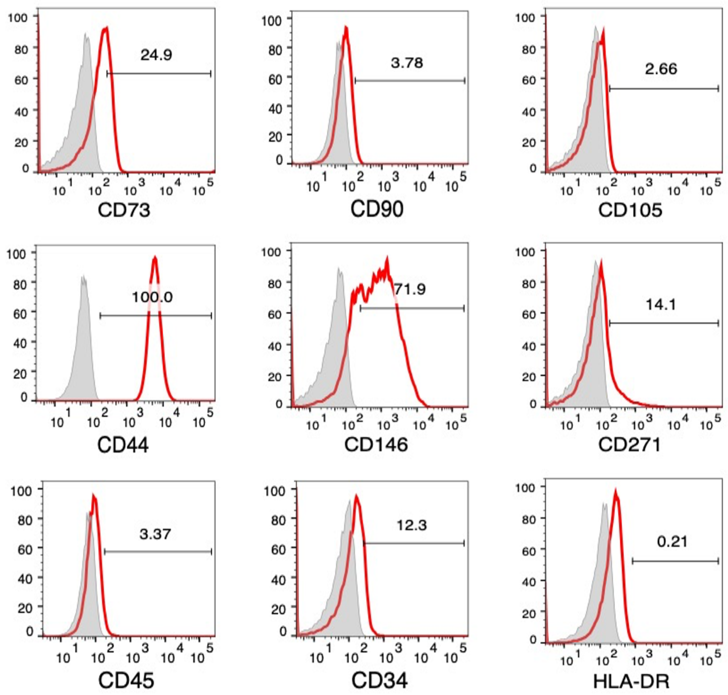


**Supplementary Figure 3.** Flow cytometry analysis of cell surface markers for oBMSC 3.

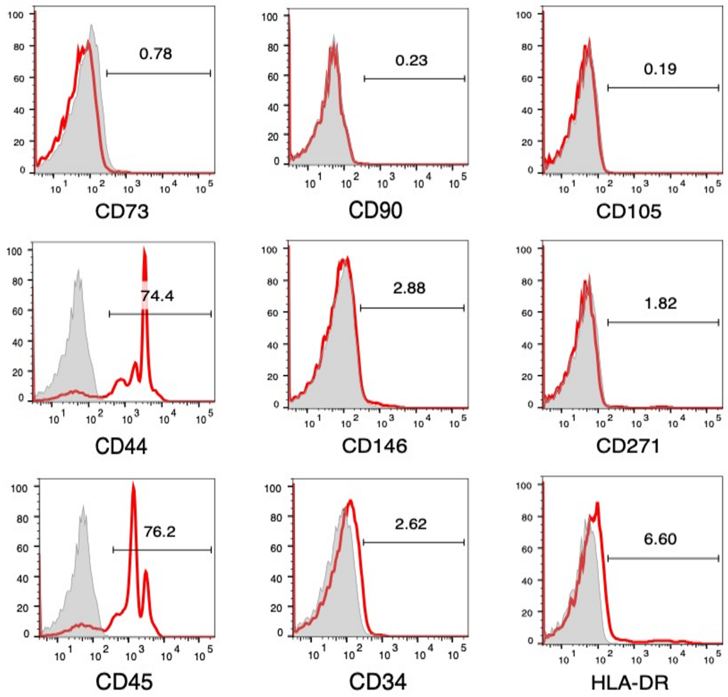


**Supplementary Figure 4.** Control flow cytometry analysis of cell surface markers for fresh sheep mononuclear cells (MNC) isolated from sheep bone marrow aspirate.

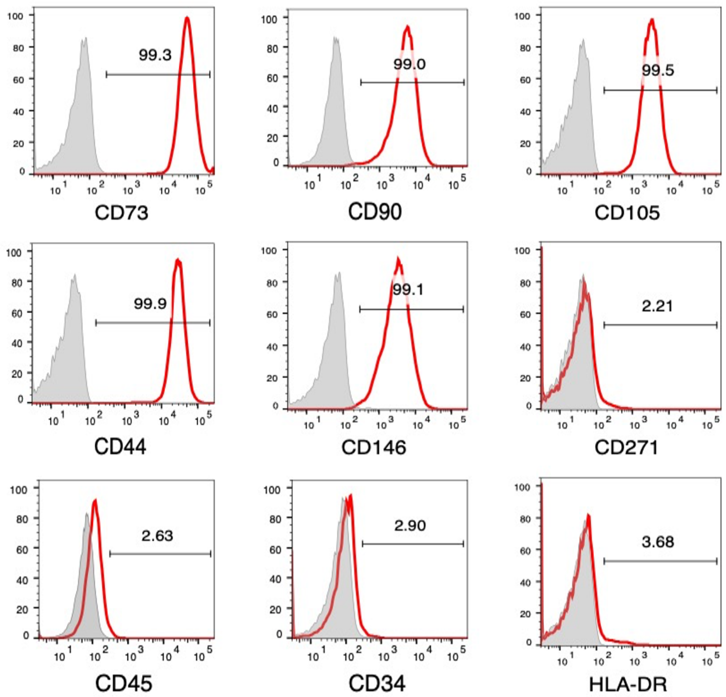


**Supplementary Figure 5.** Control flow cytometry analysis of cell surface markers on expanded hBMSC.

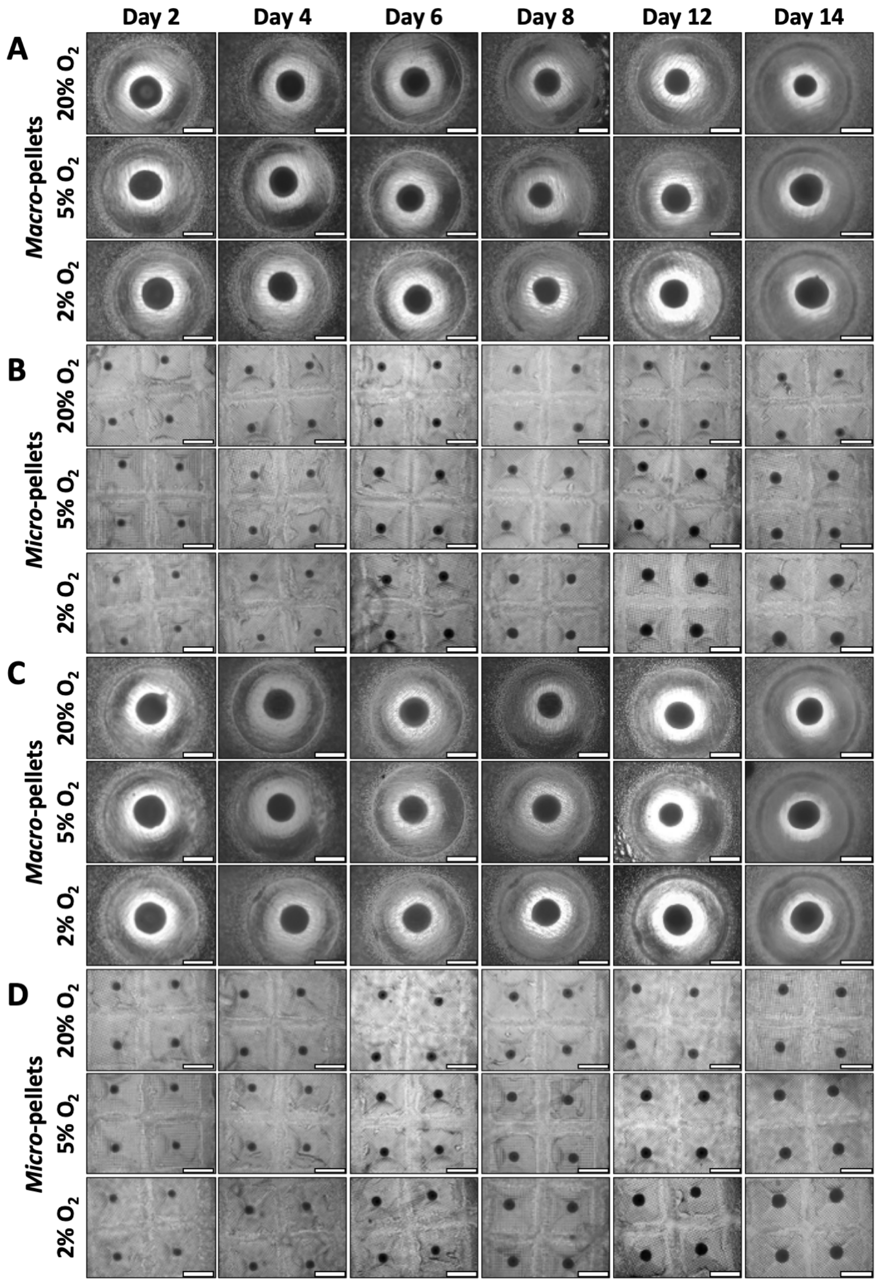


**Supplementary Figure 6.** Microscopic images over a 14-day culture period of two additional oBMSC donors cultured as *macro*-pellets in deep-well plates (A and C), and cultured as *micro*-pellets in the *Microwell-mesh* platform (B and D). Scale bar = 1 mm.

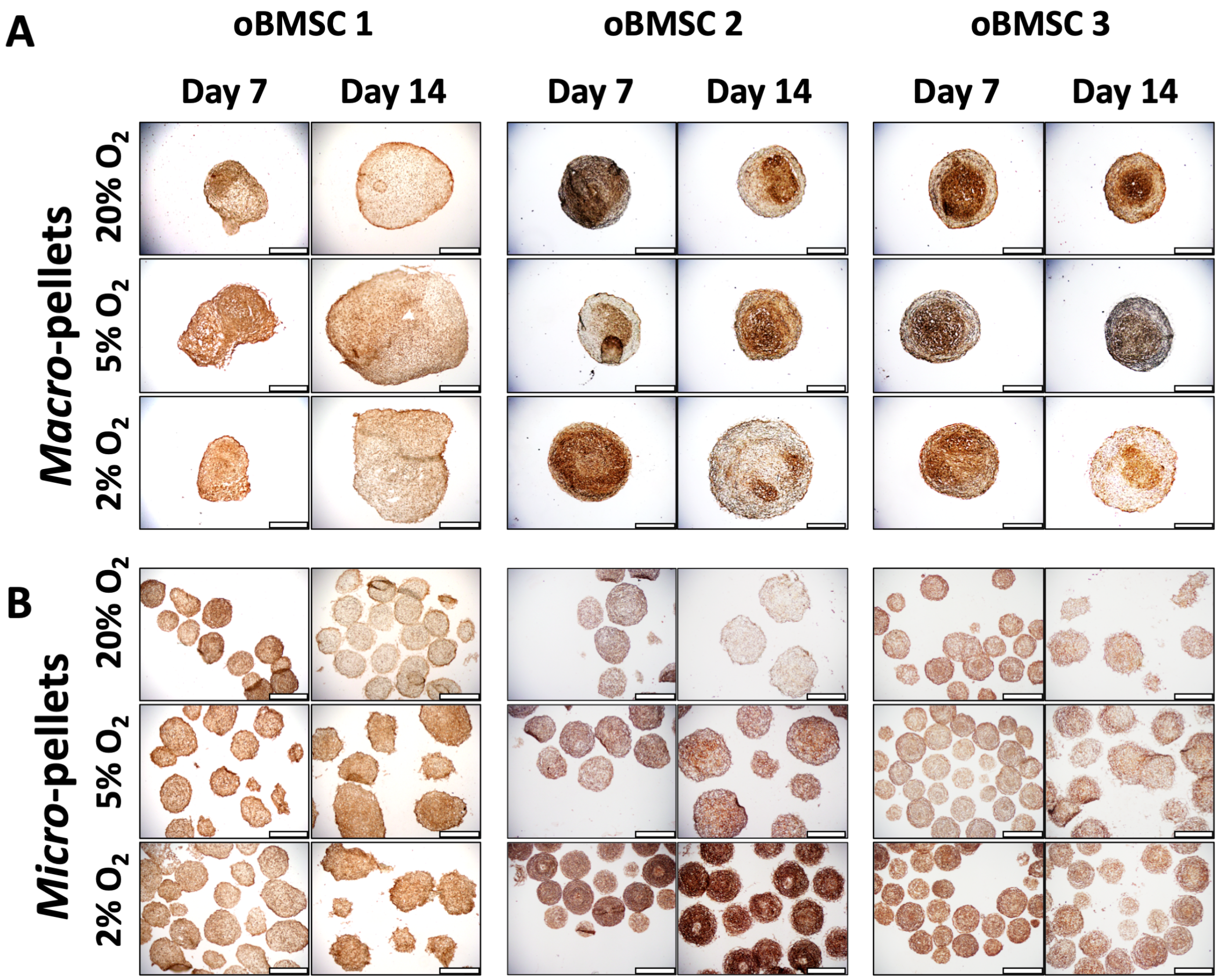


**Supplementary Figure 7.** Type II collagen staining in A) *macro*-pellet and B) *micro*-pellet sections. Scale bar = 400 µm.

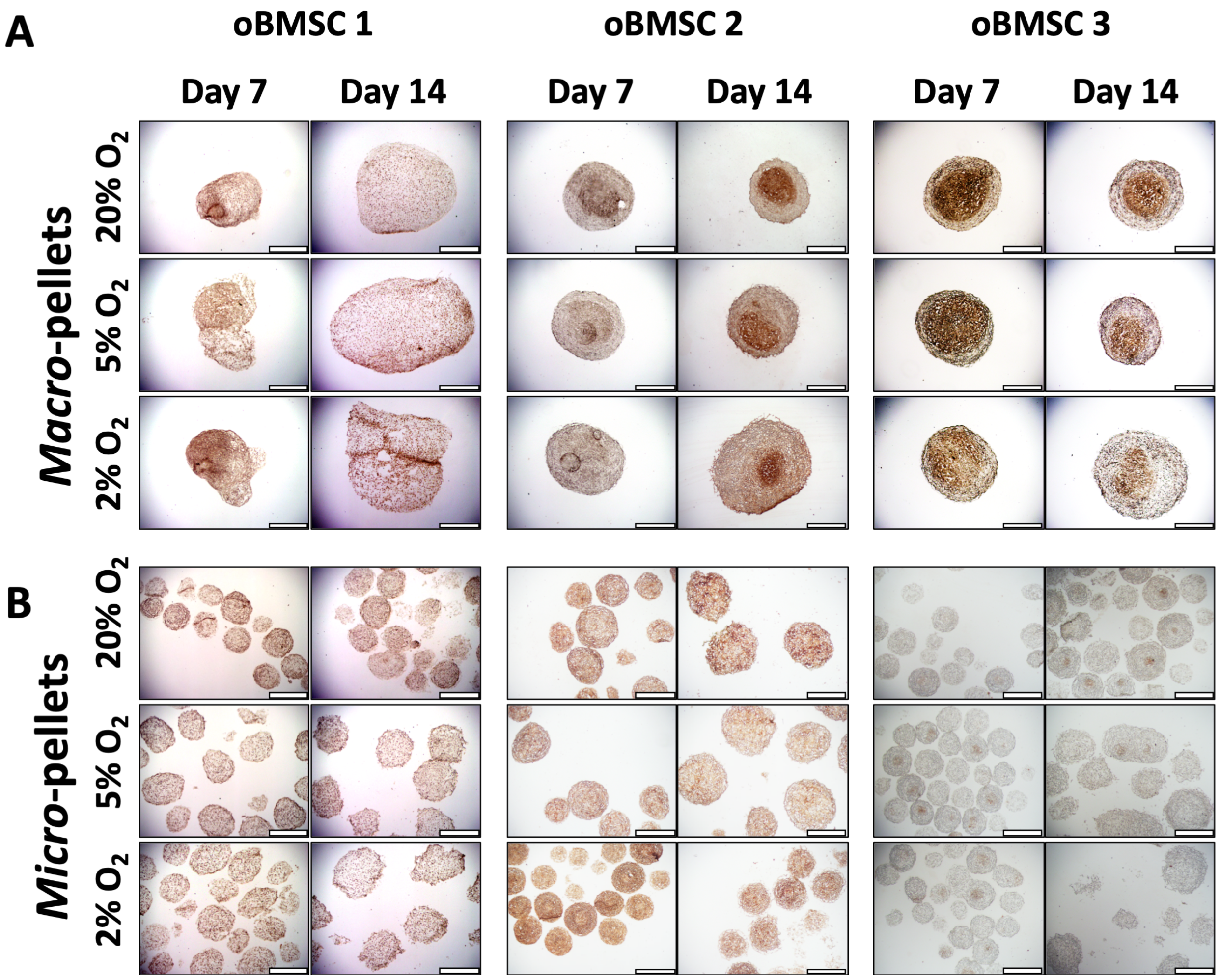


**Supplementary Figure 8.** Type I collagen staining in A) *macro*-pellet and B) *micro*-pellet sections. Scale bar = 400 µm.

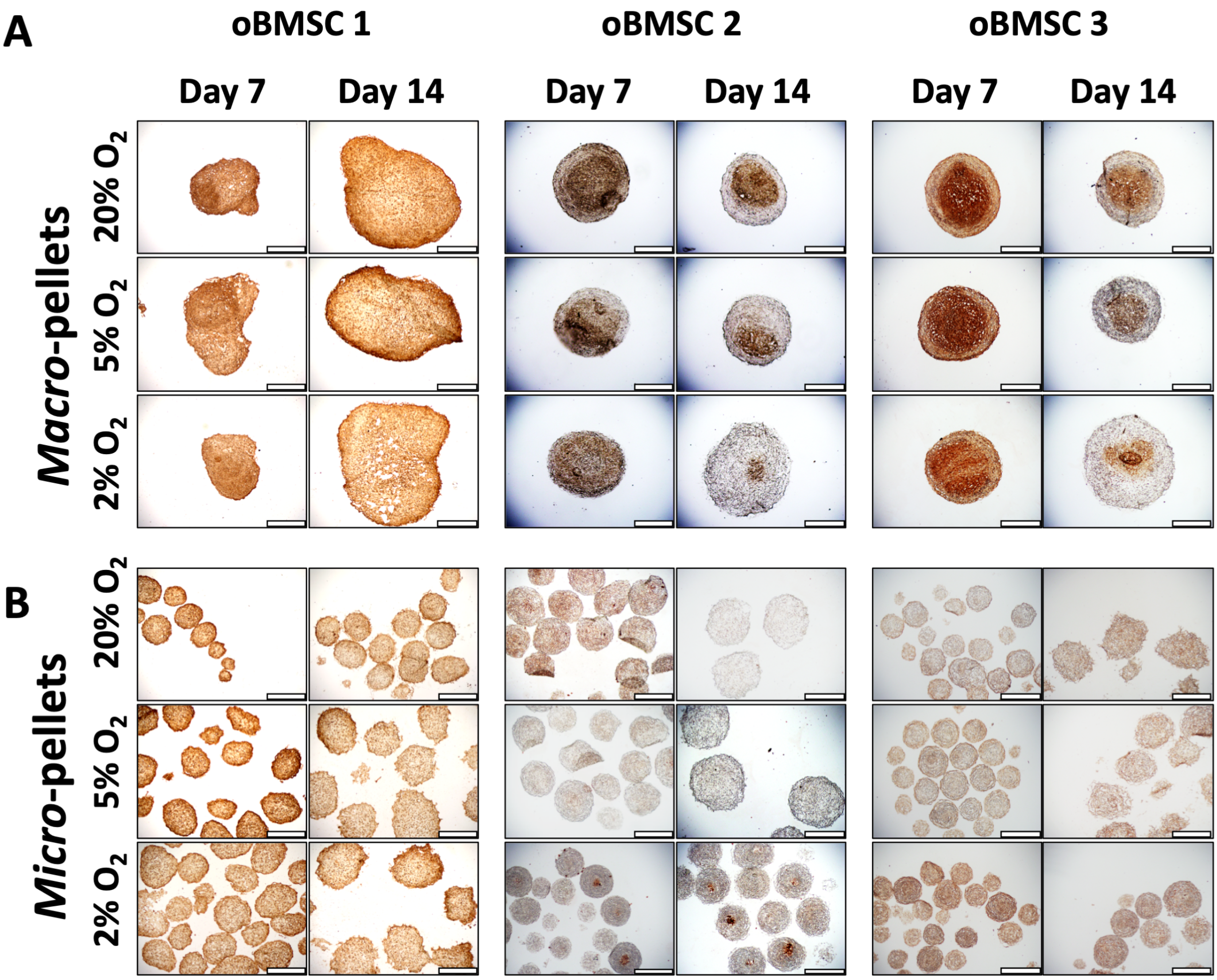


**Supplementary Figure 9.** Type X collagen staining in A) *macro*-pellet and B) *micro*-pellet sections. Scale bar = 400 μm.

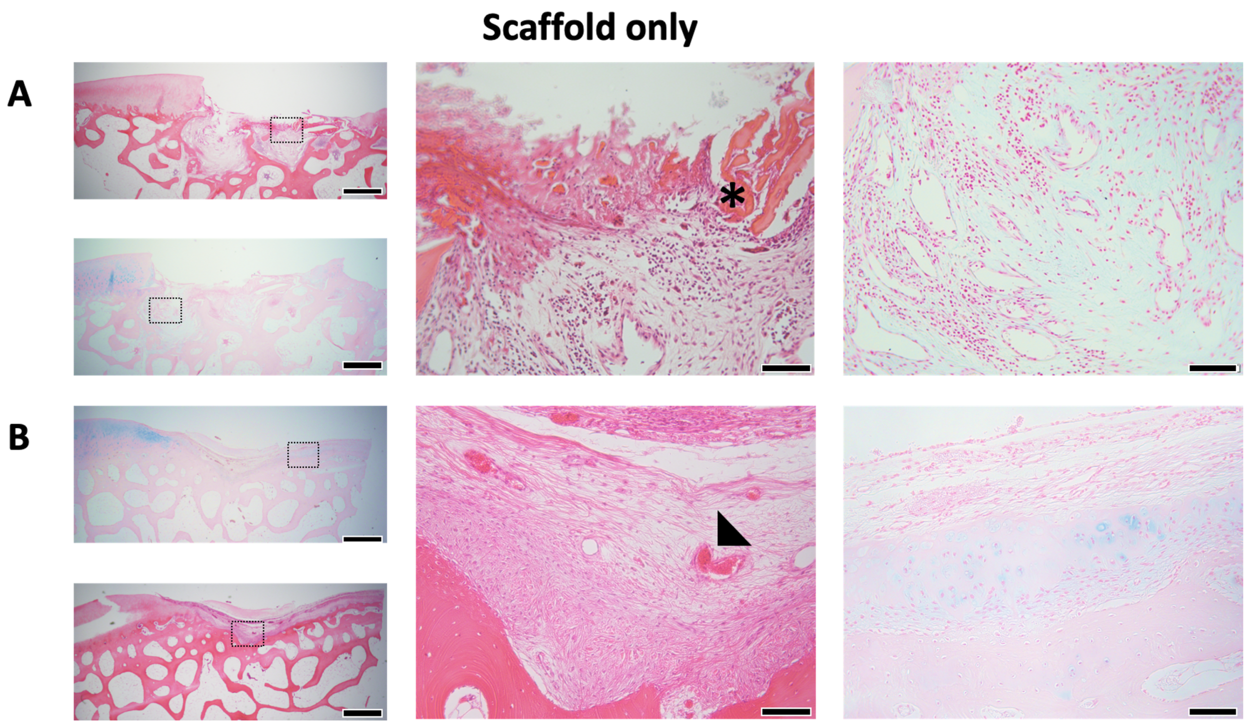


**Supplementary Figure 10.** Scaffold-only controls for A) oACh sheep pilot, and B) oBMSC sheep pilot. The middle and right images are enlarged from the boxes indicated in the images on the left. The * points to non-degraded scaffold, and the triangle points to blood vessel. Tissue was fibrous with negligible repair.

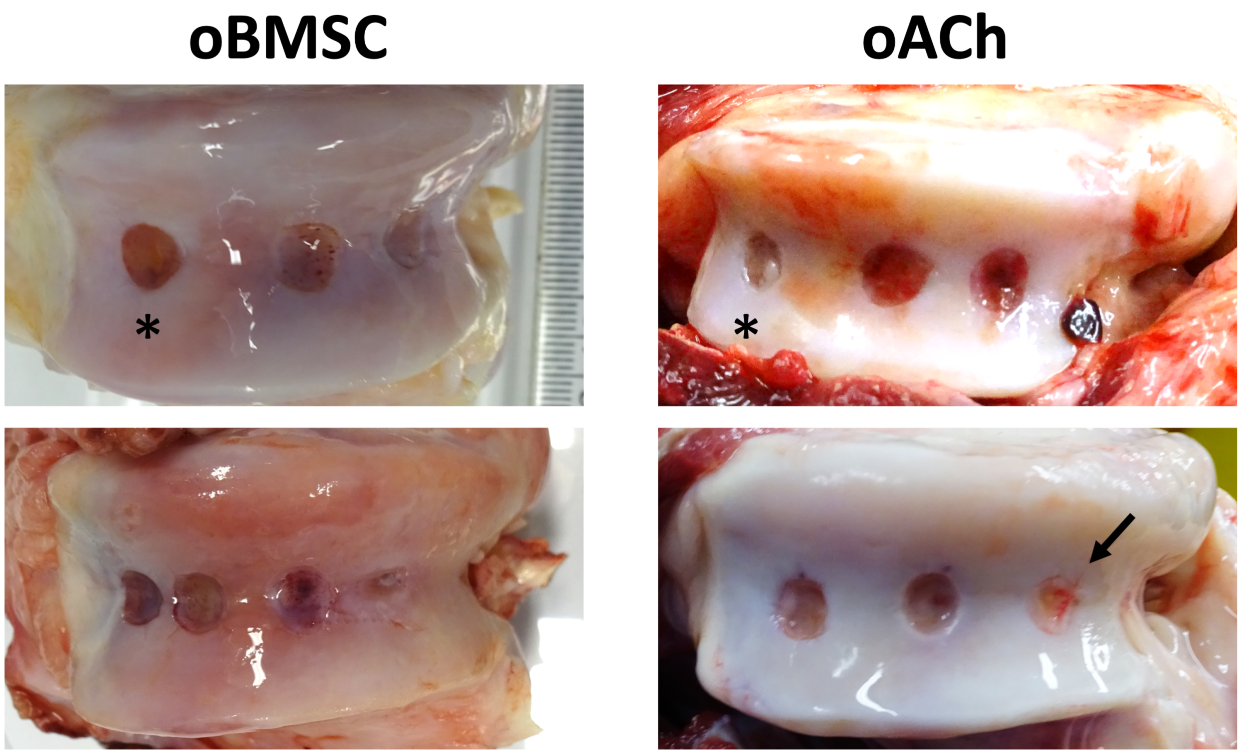


**Supplementary Figure 11.** Photos of joints 8 weeks after surgery. Asterisk indicates scaffold-only controls. Arrow points to the best repair observed with oACh *micro*-pellets.
